# A geometric shape regularity effect in the human brain

**DOI:** 10.1101/2024.03.13.584141

**Authors:** Mathias Sablé-Meyer, Lucas Benjamin, Cassandra Potier Watkins, Chenxi He, Maxence Pajot, Théo Morfoisse, Fosca Al Roumi, Stanislas Dehaene

## Abstract

The perception and production of regular geometric shapes, a characteristic trait of human cultures since prehistory, has unknown neural mechanisms. Behavioral studies suggest that humans are attuned to discrete regularities such as symmetries and parallelism, and rely on their combinations to encode regular geometric shapes in a compressed form. To identify the brain systems underlying this ability, as well as their dynamics, we collected functional MRI in both adults and six-year-olds, and magnetoencephalography data in adults, during the perception of simple shapes such as hexagons, triangles and quadrilaterals. The results revealed that geometric shapes, relative to other visual categories, induce a hypoactivation of ventral visual areas and an overactivation of the intraparietal and inferior temporal regions also involved in mathematical processing, whose activation is modulated by geometric regularity. While convolutional neural networks captured the early visual activity evoked by geometric shapes, they failed to account for subsequent dorsal parietal and prefrontal signals, which could only be captured by discrete geometric features or by bigger deep-learning models of vision. We propose that the perception of abstract geometric regularities engages an additional symbolic mode of visual perception.

## Introduction

> "In geometry, the essential is invisible to the eyes, one sees well only with the mind.”

> Emmanuel Giroux (French blind mathematician)

Long before the invention of writing, the very first detectable graphic productions of prehistoric humans were highly regular non-pictorial geometric signs such as parallel lines, zig-zags, triangular or checkered patterns (***Henshilwood et al., 2018***; ***Van der Waerden, 2012***). Human cultures throughout the world compose complex figures using simple geometrical regularities such as parallelism and symmetry in their drawings, decorative arts, tools, buildings, graphics and maps (***Tversky, 2011***). Cognitive anthropological studies suggest that, even in the absence of formal western education, humans possess intuitions of foundational geometric concepts such as points and lines and how they combine to form regular shapes (***Dehaene et al., 2006b***; ***Izard et al., 2011***). The scarce data available to date suggests that other primates, including chimpanzees, may not share the same ability to perceive and produce regular geometric shapes (***Close and Call, 2015***; ***Dehaene et al., 2022***; ***Sablé-Meyer et al., 2021***; ***Saito et al., 2014***; ***Tanaka, 2007***), though unintentional-but-regular mark-marking behavior have been reported in macaques (***Sueur, 2025***). Thus, studying the brain mechanisms that support the perception of geometric regularities may shed light on the origins of human compositionality and, ultimately, the mental language of mathematics. Here, we provide a first approach through the recording of functional MRI and magneto-encephalography signals evoked by simple geometric shapes such as triangles or squares. Our goal is to probe whether, over and above the pathways for processing the shapes of images such as faces, places or objects, the regularities of geometric shapes evoke additional activity.

The present brain-imaging research capitalizes on a series of studies of how humans perceive quadrilaterals (***Sablé-Meyer et al., 2021***). In that study, we created 11 tightly matched stimuli which were all simple, non-figurative, textureless four-sided shapes, yet varied in their geometric regularity. The most regular was the square, with four parallel sides of equal length and four identical right-angles. By progressively removing some of these features (parallelism, right-angles, equality of length, equality of angles), we created a hierarchy of quadrilaterals ranging from highly regular to completely irregular (Fig. 1A). In a variety of tasks, geometric regularity had a large effect on human behavior. For instance, for equal objective amounts of deviation, human adults and children detected a deviant shape more easily among shapes of high regularity, such as squares or rectangles (<5% errors), than among irregular quadrilaterals (>40% errors). The effect appeared as a human universal, present in preschoolers, first-graders, and adults without access to formal western math education (the Himba from Namibia), and thus seemingly independent of education and of the existence of linguistic labels for regular shapes. Strikingly, when baboons were trained to perform the same task, they showed no such geometric regularity effect.

**Fig. 1.**
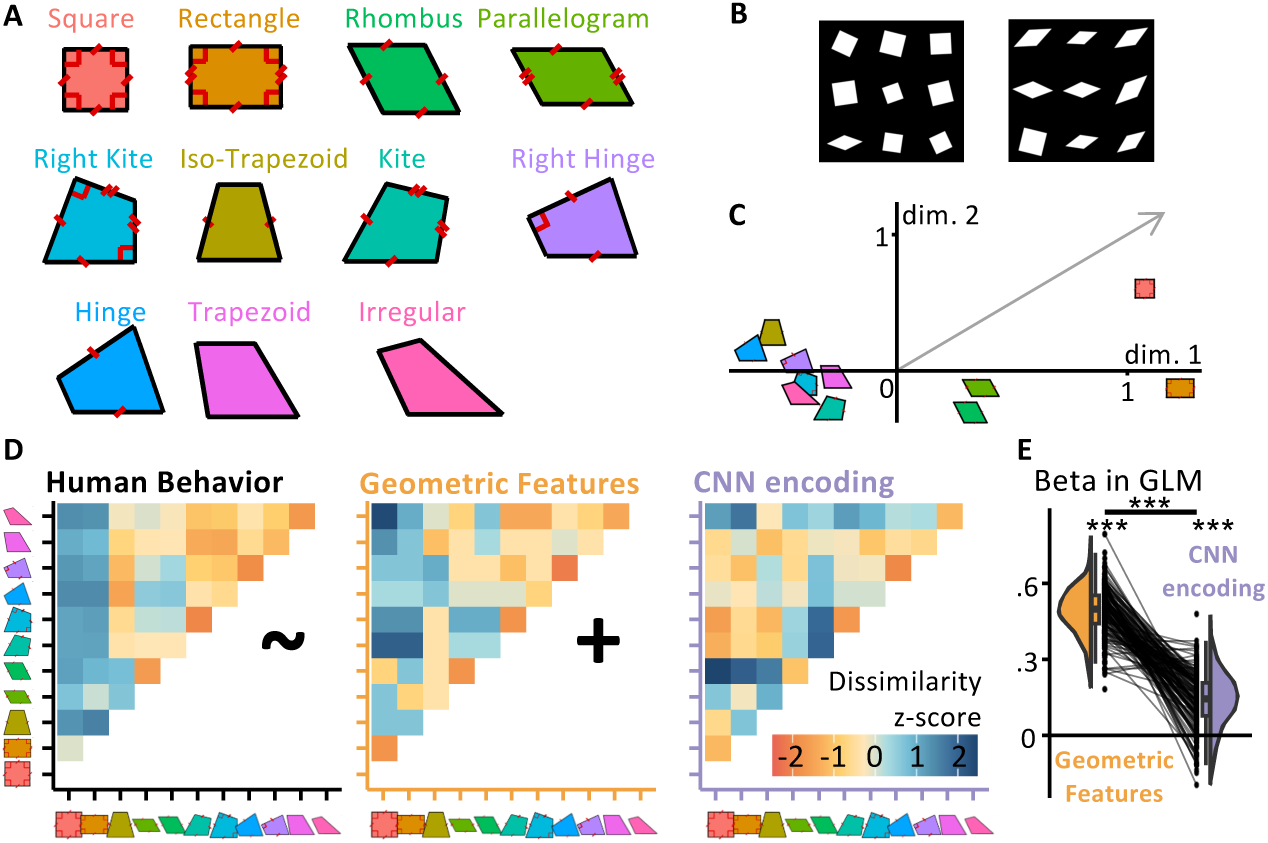
Measuring and modeling the perceptual similarity of geometric shapes. **(A)** The eleven quadrilaterals used throughout the experiments (colors are consistently used in all other figures). **(B)** Sample displays for the behavioral visual search task used to estimate the 11×11 shape similarity matrix. Participants had to locate the deviant shape. The right insert shows two trials from the behavioral visual task search task, used to estimate the 11×11 shape similarity matrix. Participants had to find the intruder within 9 shapes. **(C)** Multidimensional scaling of human dissimilarity judgments; the grey arrow indicates the projection on the MDS space of the number of geometric primitives in a shape. **(D)** The behavioral dissimilarity matrix (left) was better captured by a geometric feature coding model (middle) than by a convolutional neural network (right). The graph at right /textnf(E) shows the GLM coefficients for each participant.

Baboon behavior was accounted for by convolutional neural network (CNN) models of object recognition, but human behavior could only be explained by appealing to a representation of discrete geometric properties of parallelism, right angle and symmetry, in this and other tasks. We sometimes refer to this model as “symbolic” because it relies on discrete, exact, rule-based features rather than continuous representations (***Sablé-Meyer et al., 2022***). In this representational format, geometric shapes are postulated to be represented by symbolic expressions in a “language-of-thought”, e.g. “a square is a four-sided figure with four equal sides and four right angles” or equivalently by a computer-like program from drawing them in a Logo-like language (***Sablé-Meyer et al., 2022***).

We therefore formulated the hypothesis that, in the domain of geometry, humans deploy an additional cognitive process specifically attuned to geometric regularities. On top of the circuits for object recognition, which are largely homologous in human and non-human primates (***Bao et al., 2020***; ***Kriegeskorte et al., 2008b***; ***Tsao et al., 2008***), the human code for geometric shapes would involve a distinct “language of thought”, an encoding of discrete mathematical regularities and their combinations (***Cavanagh, 2021***; ***Dehaene et al., 2022***; ***Fodor, 1975***; ***Leeuwenberg, 1971***; ***Quilty-Dunn et al., 2022***; ***Sablé-Meyer et al., 2021***, ***2022***).

This hypothesis predicts that the most elementary geometric shapes, such as a square, are not solely processed within the ventral and dorsal visual pathways, but may also evoke a later stage of geometrical feature encoding in brain areas that were previously shown to encode arithmetic, geometric and other mathematical properties, i.e. the bilateral intraparietal, inferotemporal and dorsal prefrontal areas (***Amalric and Dehaene, 2016***, ***2019***). We hypothesized that (1) such cognitive processes encode shapes according to their discrete geometric properties including parallelism, right angles, equal lengths and equal angles; (2) the brain compresses this information when those properties are more regularly organized, and thus exhibit activity proportional to minimal description length (***Chater and Vitányi, 2003***; ***Dehaene et al., 2022***; ***Feldman, 2003***); and (3) these computations occur downstream of other visual processes, since they rely on the initial output of visual processing pathways.

Here, we assessed these spatiotemporal predictions using two complementary neuroimaging techniques (functional MRI and magnetoencephalography). We presented the same 11 quadrilaterals as in our previous research and used representational similarity analysis (***Kriegeskorte et al., 2008a***) to contrast two models for their cerebral encoding, based either on classical CNN models or on exact geometric features. In the fMRI experiment, we also collected simpler images contrasting the category of geometric shapes to other classical categories such as faces, places or tools. Furthermore, to evaluate how early the brain networks for geometric shape perception arise, we collected those fMRI data in two age groups: adults, and children in 1st grade (6-years-old, this year was selected as it marks the first-year French students receive formal instruction in mathematics). If geometric shape perception involves elementary intuitions of geometric regularity common to all humans, then the corresponding brain networks should be detectable early on.

## Experiment 1: Estimating the Representational Similarity of Quadrilaterals with Online Behavior

Our research focuses on the 11 quadrilaterals previous used in our behavioral research (***Sablé-Meyer et al., 2021***), which are tightly matched in average pairwise distance between vertices and length of the bottom side, yet differ broadly in geometric regularity (Fig.1A). The goal of our first experiment was to obtain a 11×11 matrix of behavioral dissimilarities between the 11 geometric shapes shown in Fig.1A, in order to compare it with predictions from classical visual models, embodied by CNNs, as well as geometric feature models of shape perception. To evaluate perceptual similarity in an objective manner, in experiment 1 we assessed the difficulty of visual search for one shape among rotated and scaled versions of the other (Fig.1B) (***Agrawal et al., 2019***, ***2020***). Within a grid of 9 shapes, 8 are similar and 1 is different, and participant have to click on it. Intuitively, if two shapes are very dissimilar, we expect both the response time and the error rate of finding one exemplar of one shape amongst exemplars of the other shape to be low. Conversely, we expect both to be high if shapes are similar. This gives us an empirical measure of shape dissimilarity which we can compare to the distance predicted by different models.

### Results

The 11×11 dissimilarity matrix, estimated by aggregating response time and errors from n=330 online participants is shown in Fig.1D. The distance was estimated as the average success rate divided by the average response time; method section for details. To better understand its similarity structure, we performed 2-dimensional ordinal Multi-Dimensional Scaling (MDS) projection (Fig.1C) (***De Leeuw and Mair, 2009***). The projection of the 11 shapes on the first dimension showed a strong geometric regularity, with the square and the rectangle landing at the far right, rhombus and parallelogram in the middle, and less regular shapes at the far left. Thus, human perceptual similarity seemed primarily driven by geometric properties. To quantify this resemblance using that 2D MDS projection, we examined the vector which corresponded to simply counting the number of geometric regularity (number of right angles, pairs of parallel lines, pairs of equal angles and pair of sides of equal length). This vector (shown in grey in Fig. 1C) had a projection that was significantly different from 0 for both principal axes (both p<.01).

Most diagnostically, we compared the full human dissimilarity matrix to those generated by two competing models of shape processing (***Sablé-Meyer et al., 2021***). The geometric feature model proposes that each shape is encoded by a feature vector of its discrete geometric regularities and predicts dissimilarity by counting the number of features not in common: this makes squares and rectangles very similar, but squares and irregular quadrilaterals very dissimilar. On the other hand, we operationalize our visual model by propagating shapes through a feedforward CNN model of the ventral visual pathway (we use Cornet-S, but see Fig.S1 in Supplementary Materials for other CNNs and other layers of Cornet-S, with unchanged conclusions). Shape similarity was estimated as the crossnobis distance between activation vectors in late layers (***Walther et al., 2016***). Note that these two models are not significantly correlated (r^2^=.04, p>.05).

Multiple regression of the human dissimilarity matrix with the predictions of those two visual and geometric feature models (Fig.1E) showed that both contributed to explaining perceived similarity (p*s*<.001), but that the weight of the geometric feature model was 3.6 times larger than that of the visual model, a significant difference (p<.001). This finding supports the prior proposal that the two strategies contribute to human behavior, but that the geometric feature based one dominates, especially in educated adults (***Sablé-Meyer et al., 2021***). Additional models and comparisons are presented in Fig. S5: in particular, we have included two distance measures based on skeletal representations (***Ayzenberg and Lourenco, 2019***; ***Morfoisse and Izard, 2021***), both of which performed better than chance but significantly less than the geometric feature model. Finally, to separate the effect of name availability and geometric features on behavior, we replicated our analysis after removing the square, rectangle, trapezoids, rhombus and parallelogram from our data (Fig. S5D). This left us with five shapes and an RDM with 10 entries, When regressing it in a GLM with our two models, we find that both models are still significant predictors (p<.001). The effect size of the geometric feature model is greatly reduced, yet remained significantly higher than that of the neural network model (p<.001).

## Experiment 2: fMRI Geometric Shape Localizer

To understand the neural underpinning of the cognition of geometric shape using fMRI, we started with the simplest foray into geometric shape perception by including geometric shapes as an additional visual category in a standard visual localizer used in the lab (Fig. 2). Across short mini-blocks, this fMRI run probed whole-brain responses to geometric shapes (triangles, squares, hexagons, etc.) and to a variety of matched visual categories (faces, objects, houses, arithmetic, words and Chinese characters). In 20 adults in functional MRI (9 females; 19-37 years old, mean age 24.6), we collected three localizer runs using a fast miniblock design. To maintain attention, participants were asked to detect a rare target, which could appear in any miniblock.

**Fig. 2.**
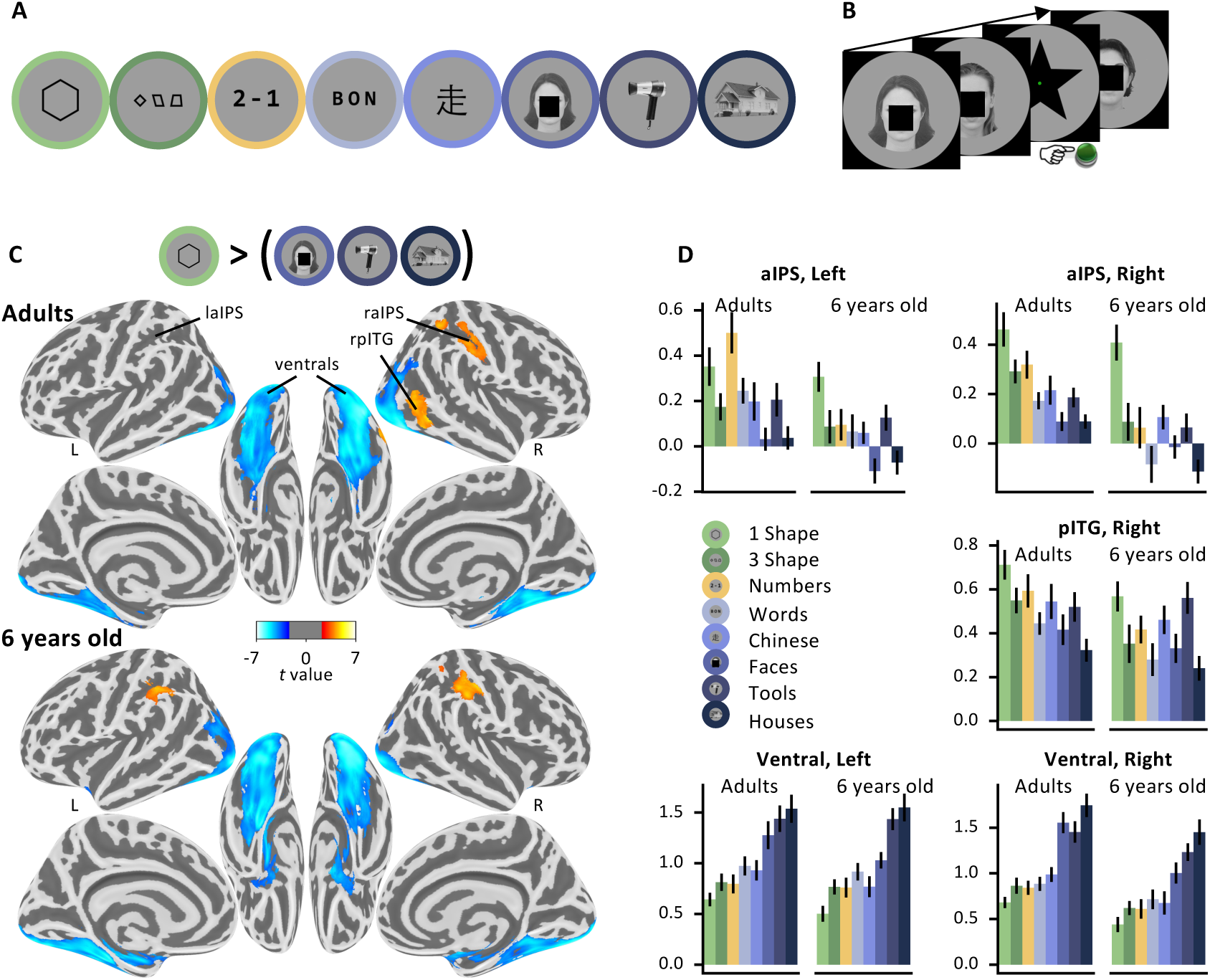
Localizing the brain systems involved in geometric shape perception. **(A)** Visual categories used in the localizer. Here and in the rest of this document, faces have been masked to comply with bioRxiv’s policy, but participants were shown unmasked faces. **(B)** Task: Passive presentation by miniblocks of consistent visual categories. In some miniblock, among a series of 6 pictures from a given category, participants had to detect a rare target star. **(C)** Statistical map associated with the contrast “single geometric shape > faces, houses and tools”, projected on an inflated brain (top: adults; bottom: children; clusters significant at cluster-corrected p<.05 with nonparametric two-tailed bootstrap test as reported in the text). **(D)** BOLD response amplitude (regression weights, arbitrary units) within each significant cluster with subject-specific localization. Geometric shapes activate the intraparietal sulcus (IPS) and posterior inferior temporal gyrus (pITG), while causing reduced activation in broad bilateral ventral areas compared to other stimuli; see Fig.S4 for analysis of subject-specific ventral subregions.

### Results

#### Reduced activity to geometric shapes in ventral visual cortex

Classical ventral visual category-specific responses, for instance to faces or words, were easily replicated (Fig.2; Fig.S2). However, when contrasting geometric shapes to faces, houses or tools, we observed a massive under-activation of bilateral ventral occipito-temporal areas (unless otherwise stated, all statistics are at voxelwise p<.001, clusterwise p<.05 permutation-test corrected for multiple comparisons across the whole brain). All of the regions specialized for words, faces, tools or houses showed this activity reduction when presented with geometric shapes (Fig.2C; Table S1).

This group analysis was further supported by subject-specific analyses of ROIs specialized for various categories of images (see Appendix and Fig.S4). First, unsurprisingly, the FFA, which is known for its strong category-selectivity, showed the lowest responses to geometric shapes in the fusiform face area. Second, in a subject-specific ROI analysis of the visual word form area (VWFA), identified by its stronger response to alphabetical stimuli than to face, tools and houses, no activity was evoked by single shapes, or strings of three shapes, above the level of other image categories such as objects or faces. This finding eliminates the possibility that geometric shapes might have been processed similarly to letters. Similarly, one could have thought that geometric shapes would be processed together with other complex man-made objects, which are often designed to be regular and symmetrical. However, geometric shapes again yielded a low activation in individual ventral visual voxels selective to tools, thus refuting this possibility. Finally, geometric shapes could have been encoded in the parahippocampal place area (PPA), which is known to encode the geometry of scenes, including abstract ones presented as Lego blocks (***Epstein et al., 1999***). However, again, geometric shapes actually induced minimal activity in individually defined PPA voxels (see Fig.S4 for a summary).

#### Increased activity to geometric shapes in intraparietal and inferior temporal cortices

While the activity of the ventral visual cortex thus seemed to be globally reduced during geometric shape perception, we observed, conversely, a superior activation to geometric shapes than to face, tools and houses in only two significant positive clusters, in the right anterior intraparietal sulcus (aIPS) and posterior inferior temporal gyrus (pITG) bordering on the lateral occipital sulcus. At a lower threshold (voxel p<.001 uncorrected), the symmetrical aIPS in the left hemisphere was also activated (also see Supplementary Materials for additional results concerning the “3-shapes” condition and with both shape conditions together).

Those areas are similar to those active during number perception, arithmetic, geometric sequences, and the processing of high-level math concepts (***Dehaene et al., 2022***; ***Amalric and Dehaene, 2016***, ***2019***). To test this idea formally, we used the localizer to identify ROIs activated by numbers more than words. This contrast identified a left IPS cluster (p<.05), while the symmetrically identified cluster in right IPS did not reach significance at the whole brain level (p=.18). In both cases, however, the ROI was also significantly more activated by geometric shapes than other visual categories.

The observed overlap with number-related areas of the IPS is compatible with our hypothesis that geometric shapes are encoded as mental expressions that combines number, length, angle and other geometric features. However, the association between geometric shapes and other arithmetic or mathematical properties could be acquired during schooling. To test whether the brain activity observed in adults reflects a basic intuition of geometry which is also present early on in child development, we replicated our fMRI study in 22 6-year-old first graders. When comparing the single shape condition and faces, houses and tools, we observed the same reduction in ventral visual activity. We also observed greater aIPS activity, now significant in both hemispheres-though the identifed right pITG cluster did not reach significance. Still, the right pITG voxels extracted from adults reached significance in children for the shape versus face, houses and tools. Conversely, the left aIPS voxels extracted in children reached significance in adults. Indeed, the activation profiles were quite similar in both age groups (Fig.2D; see Fig.S3A and Fig.S3B for uncorrected statistical maps of adults and children, which are quite similar). In particular, aIPS activity was strongest to geometric shapes in children, while in adults strong responses to both geometric shapes and numbers were seen, indicating an overlap with previously observed areas involved in arithmetic (***Amalric and Dehaene, 2019***; ***Pinheiro-Chagas et al., 2018***).

To better establish the correspondence between our findings and previous work, we evaluated the “geometric shape > other visual categories” contrast in six ROIs previously identified as part of the math-responsive network (***Amalric and Dehaene, 2016***): bilateral IPS, bilateral MFG, and bilateral pITG. In all ROIs, in both adults and children, the average geometric shape contrast was significant at the p<.05 level, except for the left posterior ITG in children (p=.09). Nevertheless, cautiousness is required here because activation overlap could arise without indicating that the same exact circuits and processes are involved especially in smoothed group-level images. To partially mitigate this problem and test for subject-level overlap between the activations to geometric shapes and to numbers, we turned to within-subject analyses. We assessed whether the patterns of activation evoked by numbers and geometric shapes were more similar to each other than those evoked by numbers and other categories. For each subject, we computed the crossnobis distance between the activations evoked by each of the experimental conditions. We then compared the distances for numbers versus geometric shapes, and for numbers versus the average of the other categories. We found that, in adults, in four of our five ROIs, the activations to geometric shapes were indeed more similar to numbers than to other categories (p<.05 in lIPS, rITG, and bilateral ventral ROIs, p=.17 in rIPS). In children, the pattern of similarity were too noisy to be conclusive for bilateral IPS or the ITG, but the finding remained significant in bilateral ventral areas (both p<.05).

In sum, geometric shapes led to reduced activity in ventral visual cortex, where other categories of visual images show strong category-specific activity. Instead, geometric shapes activated areas independently found to be activated by math- and number-related tasks, in particular the right anterior intraparietal sulcus.

## Experiment 3: fMRI of the geometric intruder task

In a second fMRI experiment, we measured the detailed fMRI patterns evoked by our quadrilaterals, with the goal to submit them to a representational similarity analysis and test our hypothesis of a double dissociation between regions encoding visual (CNN) versus geometric codes. Adults and children performed an intruder task similar to our previous behavioral study (***Sablé-Meyer et al., 2021***), with miniblocks allowing us to evaluate the activity pattern evoked by each quadrilateral shape. To render the task doable by 6-year-olds, we tested only 6 quadrilaterals, displayed as two half-circles of three items, and merely asked participants whether the intruder was on the left or the right of the screen (Fig.3).

**Fig. 3.**
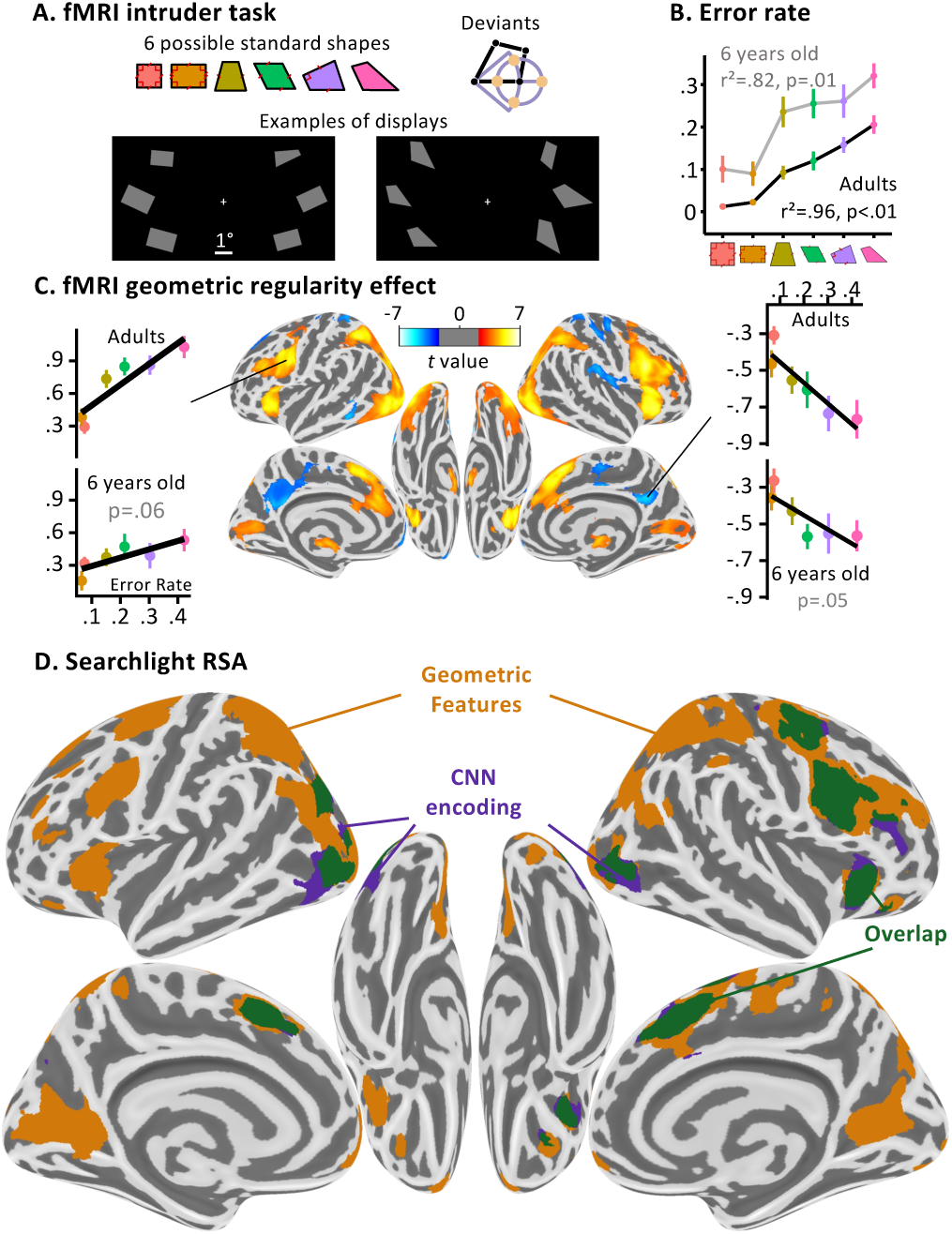
Dissociating two neural pathways for the perception of geometric shape. **(A)** fMRI intruder task. Participants indicated the location of a deviant shape via button clicks (left or right). Deviants were generated by moving a corner by a fixed amount in four different directions. **(B)** Performance inside the fMRI: both populations tested displayed an increasing error rate with geometric shape complexity, which significantly correlates with previous data collected online. **(C)** Whole brain correlation of the BOLD signal with geometric regularity in adults, as measured by the error rate in a previous online intruder detection task (***Sablé-Meyer et al., 2021***). Positive correlations are shown in red and negative ones in blue. Voxel threshold p<.001, cluster-corrected by permutation at p<.05. Side panels show the activation in two significant ROIs whose coordinates were identified in adults and where the correlation was also found in children (one-tailed test, corrected for the number of ROIs tested this way). **(D)** Whole-brain searchlight-based RSA analysis in adults (same statistical thresholds). Colors indicate the model which elicited the cluster: purple for CNN encoding, orange for the geometric feature model, green for their overlap.

### Results

Behavioral performance inside the scanner replicated the geometric regularity effect in adults and children: performance was best for squares and rectangles and decreased linearly for figures with fewer geometric regularities (Fig.3B). Behavior in the fMRI scanner was significantly correlated with an empirical measure of geometric regularity measured with an online intruder detection task (***Sablé-Meyer et al., 2021***), which is used in all following analyses whenever we refer to geometric regularity. The neural bases of this effect were studied at the whole-brain level by searching for areas whose activation varied monotonically with geometrical regularity. In adults, this contrast identified a broad bilateral dorsal network including occipito-parietal, middle frontal, anterior insula and anterior cingulate cortices, with accompanying deactivation in posterior cingulate (Fig.3C). This broad network, however, probably encompassed both regions involved in geometric shape encoding and those involved in the decision about the intruder task, whose difficulty increased when geometric regularity decreased. To isolate the regions specifically involved in shape coding, we performed searchlight RSA analyses, which focused on the pattern rather than the level of activation. Within spheres spanning the entire cortex, we asked in which regions the similarity matrix between the activation patterns evoked by the six shapes was predicted by the CNN encoding model, the geometric feature model, or both (Fig.3D; Table S2). In adults, the CNN encoding model predicted neural similarity in bilateral lateral occipital clusters, while the geometric feature model yielded a large set of clusters in occipito-parietal, superior prefrontal and right dorsolateral prefrontal cortex. Calcarine cortex was also engaged, possibly due to top-down feedback (***Williams et al., 2008***). Both CNN encoding and geometric feature models had overlapping voxels in anterior cingulate and right premotor cortex, possibly reflecting the pooling of both visual codes for decision-making.

In children, possibly because the task difficulty was high, few results were obtained. The correlation of brain activity with regularity at the whole-brain level yielded a single significant cluster (see Figure S3C for an uncorrected statistical map (voxelwise p<.001), with the single significant whole-brain, cluster corrected significant cluster (p=.012) in the ventromedial prefrontal cortex). When testing the ROIs from the adults in children, after correcting for multiple comparisons across the 14 ROIs, only the posterior cingulate was significant (p=.049), though two clusters were close to significance, both with p=.06: one positive in the left precentral gyrus (shown in Fig.3C), and one negative in the dorsolateral prefrontal cortex. Testing the ROIs identified in the visual localizer showed that they all exhibited a positive geometric difficulty effect in adults (all p<.05), but did not reach significance in children, possibly due to excessive noise (as indicated by the much higher error rate in the task inside the scanner). In the searchlight analysis, no cluster associated to the geometric feature model reached significance at the whole-brain level (Fig.S3D).

However, a right lateral occipital cluster was significantly captured by the CNN encoding model in children (p=.019) and its symmetrical counterpart was close to the significance threshold (p=.062) (Fig.S3D). This result might indicate that geometric features are not well differentiated prior to schooling. It could also reflect that children weight the geometric feature strategy less, as was found in previous work (***Sablé-Meyer et al., 2021***); combined with difficulty of obtaining precise subject-specific activation patterns in young children, this could make the geometric feature strategy harder to localize.

## Experiment 4: oddball paradigm of geometric shapes in MEG

The temporal resolution of fMRI does not allow to track the dynamic of mental representations over time. Furthermore, the previous fMRI experiment suffered from several limitations. First, we studied six quadrilaterals only, compared to 11 in our previous behavioral work (***Sablé-Meyer et al., 2021***). Second, we used an explicit intruder detection, which implies that the geometric regularity effect was correlated with task difficulty, and we cannot exclude that this factor alone explains some of the activations in figure 3C (although it is much less clear how task difficulty alone would explain the RSA results in figure 3D). Third, the long display duration, which was necessary for good task performance especially in children, afforded the possibility of eye movements, which were not monitored inside the 3T scanner and again could have affected the activations in figure 3C.

To overcome those issues, we replicated the experiment in adult magnetoencephalography (MEG) with three important changes; (1) all 11 quadrilaterals were studied; (2) participants were simply asked to fixate and attend to every shape, without performing any explicit task; (3) shapes were presented serially, one at a time, at the center of screen, with small random changes in rotation and scaling parameters; (4) In miniblocks of 30 seconds each, a fixed quadrilateral shape appeared repeatedly, interspersed with rare intruders whose bottom right corner was shifted by a fixed amount (***Sablé-Meyer et al., 2021***). This design allowed us to study the neural mechanisms of the geometric regularity effect without confounding effects of task, task difficulty, or eye movements. Would the shapes be automatically encoded according to their geometric regularities even in such a passive context? And would brain responses indicate that the intruders continued to be detected more easily among geometric regular shapes than among irregular ones, as previously found behaviorally in an active task (***Sablé-Meyer et al., 2021***)?

### Results

In spite of the passive design, MEG signals revealed an automatic detection of intruders, driven by geometric regularity. We trained logistic regression decoders to classify the MEG signals at each time point following a shape as arising from a reference shape or from an intruder (see Method and Fig.4A). Overall, the decoder performed above chance level, reaching a peak at 428ms, indicating the presence of brain responses specific to intruders. Crucially, although trained on all shapes together, the decoder performed better with geometrically regular shapes than with irregular shapes (and better than chance for each), indicating that oddball shapes were more easily detected within blocks of regular shapes, as previously found behaviorally (***Sablé-Meyer et al., 2021***). Indeed, a regression of decoding performance on geometrical regularity yielded a significant spatio-temporal cluster of activity, which first became significant around ∼160ms and peaked at 432ms (here and after, temporal clusters are purposefully reported with approximate bounds following (***Sassenhagen and Draschkow, 2019***)). A geometrical regularity effect was also seen in the latency of the outlier response (See Fig.4A; correlation between geometric regularity and the latency when average decoding performance first exceeded 57% correct; one-tailed t-test of each participant’s regression slope against 0: t=-1.83, p=.041) indicating that oddballs yielded both a larger and a faster response when the shapes were geometrically more regular. The same effect was found when training separate outlier decoders for each shape, thus refuting an alternative hypothesis according to which all outliers evoke identical amounts of surprise, but in different directions of neural space (Fig.4B). Overall, the results fully replicate prior behavioral observations of the geometric regularity effect (***Sablé-Meyer et al., 2021***) and suggest that the computation of a geometric code and its deviations occurs under passive instructions and starts with a latency of about ∼150ms.

**Fig. 4.**
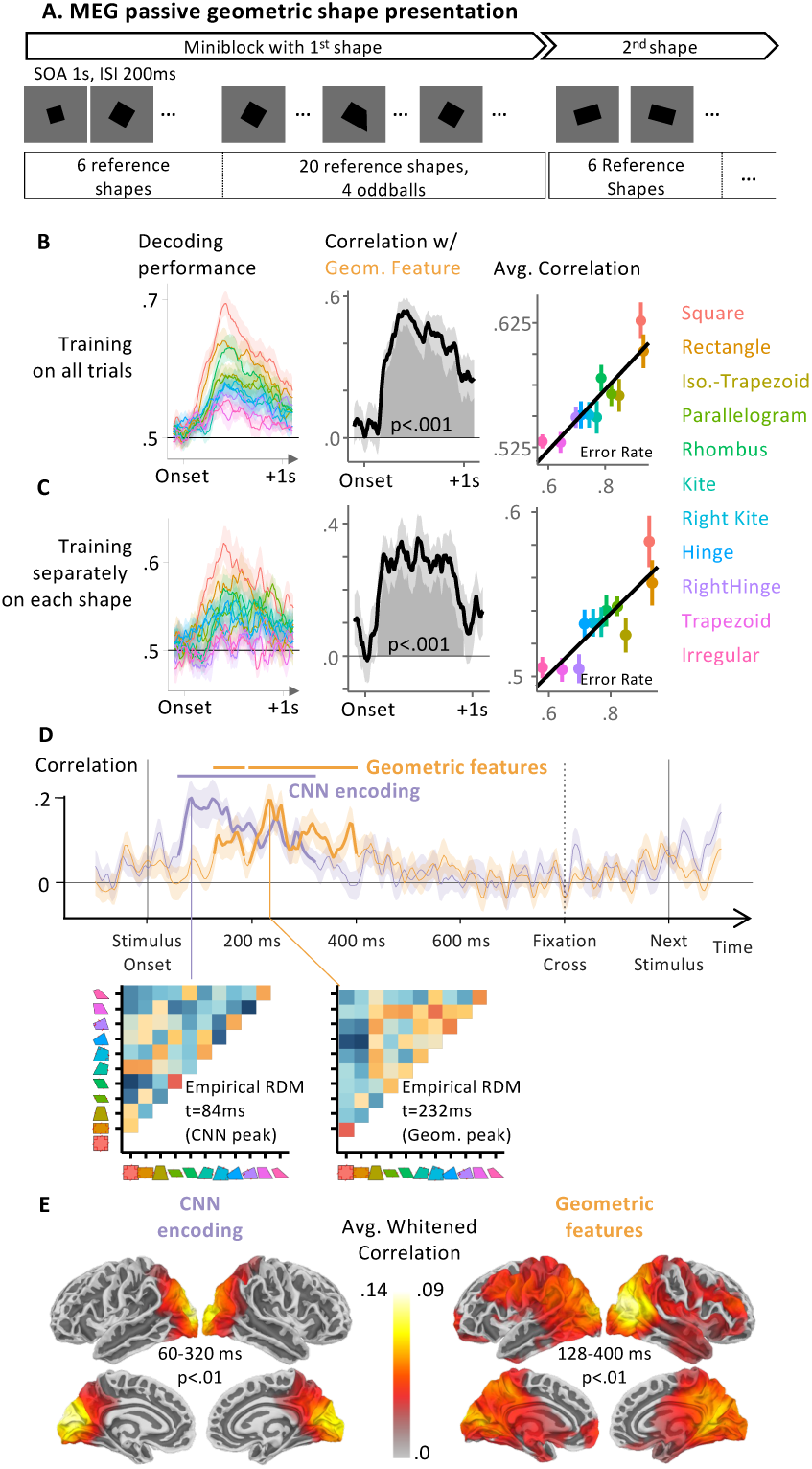
Using MEG to time the two successive neural codes for geometric shapes. **(A)** Task structure: participants passively watch a constant stream of geometric shapes, one per second (presentation time 800ms). The stimuli are presented in blocks of 30 identical shapes up to scaling and rotation, with 4 occasional deviant shape. Participants do not have a task to perform beside fixating. **(B,C)** Performance of a classifier using MEG signals to predict whether the stimulus is a regular shape or an oddball. Left: performance for each shape; middle: correlation with geometrical regularity (same x axis as in figure 3C); right: visualization of the average decoding performance over the cluster. In B, training of the classifier was performed on MEG signals from all 11 shapes; In C, eleven different classifiers were trained separately, one for each shape. **(D)** Sensor-level temporal RSA analysis. At each time point, the 11×11 dissimilarity matrix of MEG signals was modeled by the two model RDMs in Fig. 1D, and the graph shows the time course of the corresponding whitened correlation coefficients. Below the timecourses, we display the average empirical dissimilarity matrix across participants at two notable timepoints: when the correlation with the CNN and Geometric Feature models are maximal (CNN: t=84ms; Geometric Features: 232ms) **(E)** Source-level temporal-spatial searchlight RSA. Same analysis as in C, but now after reconstruction of cortical source activity.

We next used temporal RSA to further probe the dynamics of the perception of the reference, non-intruder shapes. For each time point, we estimated the 11×11 neural dissimilarity matrix across all pairs of reference shapes using sensor data with crossnobis distances (***Walther et al., 2016***), and entered them in a multiple regression with those predicted by CNN encoding and geometric feature models. The coefficients associated to each predictor are shown in Fig. 4C. An early cluster (observed cluster extent at approximately 60 to 320ms, peak at 84ms; p<.001) showed a significant correlation of the CNN encoding model on brain similarity. It was followed by two significant cluster associated with the geometric feature model separated by two timepoints that did not pass the cluster-formation threshold (∼128-184ms then ∼196-400ms, overall peak at 232 ms; p<.001). Those results are compatible with our hypothesis of two distinct stages in geometric shape perception and suggest that a geometric feature base encoding is present, especially around ∼200-250ms. This analysis yielded similar clusters when performed on a subset of shapes that do not have an obvious name in English, as was the case for the behavior analysis (CNN Encoding: 64.0ms to 172.0ms; then 192.0ms to 296.0ms; both p<.001: Geometric Features: 312.0ms to 364.0ms with p=.008, the later timing could indicate that geometric features for less regular shapes take longer to estimate, replicating the latency effect found in the oddball decoding analysis).

To understand which brain areas generated these distinct patterns of activations, and probe whether they fit with our previous fMRI results, we performed a source reconstruction of our data. We projected the sensor activity onto each participant’s cortical surfaces estimated from T1-images. The projection was performed using eLORETA (***Jatoi et al., 2014***) and emptyroom recordings acquired on the same day to estimate noise covariance, with the default parameters of mne-bids-pipeline. Sources were spaced using a recursively subdivided octahedron (oct5). Group statistics were performed after alignement to fsaverage. We then replicated the RSA analysis with search-lights sweeping across cortical patches of 2 cm geodesic radius (Fig. 4D). The CNN encoding model captured brain activity in a bilateral occipital and posterior parietal cluster, while the geometric model accounted for subsequent activity starting at ∼200 ms and spanning over broader dorsal occipito-parietal and intraparietal, prefrontal, anterior and posterior cingulate cortices. This double dissociation closely paralleled fMRI results (compare Fig. 4D and Fig. 3D; see also Video 1 for a movie of the significant sources associated to either model across time), with greater spatial spread due to the unavoidable imprecision of MEG source reconstruction.

## Modelling of the results with alternative models of visual perception

In the above analyses, we contrasted two models for our data: a CNN versus a list of geometric properties. However, there are many other models of vision. In this section we model our data with two additional classes of models, based on the shape skeleton and principal axis theory, or vision transformers neural networks.

### Skeleton models

Skeletal representations (***Blum, 1973***) offer biologically plausible and computationally tractable models of object representation in human cognition. Several methods to derive skeletal representation from object contour have been put forward, including Bayesian estimation that trade off accuracy for conciseness (***Feldman and Singh, 2006***). More recently, distance estimators between shapes based on their skeletal structure have been used to model human behavior (***Ayzenberg et al., 2022a***; ***Ayzenberg and Lourenco, 2019***; ***Denisova et al., 2016***; ***Lowet et al., 2018***; ***Morfoisse and Izard, 2021***), and hierarchical object decomposition have been rigorously described and tested (***Froyen et al., 2015***). Remarkably, even when simply asked to tap a geometric shape on a touch-screen anywhere they want, human taps cluster along the shape skeleton (***Firestone and Scholl, 2014***).

To model our data, we needed a distance metric between two shape skeletons. We explored two metrics: one that measures the shortest distance between two skeletons after optimal alignment and rotation (***Ayzenberg and Lourenco, 2019***; ***Jacob et al., 2021***), and one that measures the difference in angle and length of matched sub-parts of the shapes’ skeleton (***Morfoisse and Izard, 2021***). Our results are summarized in Fig. S5A; the first observation is that over our 11 quadrilaterals, these two skeletal metrics were not significantly correlated, suggesting that they capture very different properties when applied to minimal geometric shapes and are possibly not best suited to characterize them. Additionally, neither of these metrics predicted the behavioral data better than the CNN models.

When tested on fMRI data, the first method based on optimal alignment and rotation did not elicit any significant cluster in either population. In adults, but not in children, the second method based on the structure of the skeletons significantly correlated with bilateral clusters in the visual cortex, as well as a significant cluster in the right premotor cortex (see Fig.S6). These areas constitute a subset of the areas identified with the CNN encoding model.

When tested on MEG data, the first method similarly did not elicit any significant temporal cluster at the p<.05 level. The second method yielded two significant temporal clusters, the first from ∼75ms to ∼260ms and the second from ∼285ms to ∼370ms. When looking at the reconstructed sources, the second temporal cluster did not yield and significant spatial cluster, and the first cluster elicited a bilateral cluster encompassing both early occipital area and the early dorsal visual stream, with no significant localization in the frontal cortex.

Overall, for our dataset, these models appear partially similar to the CNN model insofar as they capture subsets of the timings and localizations of the neural data captured by the CNN model, but they do not strongly predict the behavior data as the geometric feature model does, nor capture the broad cortical networks associated with the geometric feature model.

### Vision Transformers and Larger Neural Networks

While small CNN models have well-known limits in capturing several aspects of human perception (***Bowers et al., 2022***; ***Jacob et al., 2021***), those limits are constantly being pushed. First, connectionist models based on the more advanced transformer architecture may provide a closer match to human data, especially when it comes to symbolic or mathematical regularities (***Campbell et al., 2024***). Second, even within the CNN-like architecture, much larger neural networks, trained on vastly richer datasets, may achieve superior similarity to thumans than CORnet. We briefly consider both possibilities.

First, using the same method as with CNNs, we modeled our empirical RDMs with distance measured between pairs of shapes in the late layers of a visual transformer, DINOv2 (***Oquab et al., 2023***). In Fig. S5B, we report the empirical RDMs, the DINO RDM, as well as the CORnet CNN RDM, together with a plot of each subject’s correlation with the CNN and DINO matrices. The DINO predictor yielded a dissimilarity matrix quite similar to the human behavioral one; indeed, the distribution of coefficients for the DINO predictor was significantly greater than 0 (95% confidence interval [0.73, 0.75]; p<.001) and significantly different from the CNN predictor (p<.001). Note that in this analysis, the distribution of CNN predictors is significantly lower than zero (p<.001), which was not the case when contrasted with the symbolic model. This suggests that the DINO predictor may involve a mixture of symbolic-like features and perceptual features. Indeed, its correlation with the symbolic model is .78 and the correlation with the cornet is .56. This may explain why human are best predicted with a mixture of these two models, but with different weights: to achieve the best fit, a model with both DINO and CNN puts a high weight on DINO and then negatively corrects the excess perceptual features by putting a negative weight on the CNN model.

This analysis was confirmed by a fit of the fMRI data (Fig.S6) and MEG data (Fig.S5C): in both cases, the RSA analysis with DINO last layer yielded results that were very similar to a mixture of the CORnet CNN and the symbolic model. In fMRI in adults, a distributed set of cortical areas were significantly correlated with the DINO model, and these areas overlapped with the union of the areas predicted by both of our original models. In children, DINOv2 elicited a significant cluster in the right visual cortex, in areas overlapping with the areas activated by the CNN model in children. In MEG, a large and significant cluster was predicted by the DINO model. Its timing (significant cluster from ∼64ms to ∼400ms) overlapped with the spatiotemporal clusters found with both the CNN model and the symbolic model. Source analysis indicated that a correlation with DINO originated from widespread sources that, again, overlapped with both those found with the CNN model (early occipital areas) and the symbolic model (widespread regions included dorsal, anterior temporal, and frontal sources).

Taken together, these results suggest that the last layer of DINO is driven by both early visual and higher-level geometric features, and that both are needed to fit human data. While these findings may suggest that network architecture may be crucial to mimic human geometric perception, continuing work in our lab challenges this conclusion because similar results were obtained by performing the same analysis, not only with another vision transformer network, ViT, but crucially using a much larger convolutional neural network, ConvNeXT, which comprises ∼800 million parameters and has been trained on two billion images, likely including many geometric shapes and human drawings. For the sake of completeness, RSA analysis in sensor space of the MEG data with these two models is provided in Fig. S6. A systematic investigation of the impact of network architecture, dataset size and dataset content is beyond the scope of the present paper, but the present data could serve as a reference dataset with which to understand which of these factors ultimately cause a neural network to display symbolic properties close to those found in the human brain.

## Discussion

Our previous behavioral work suggested that the human perception of geometric shapes cannot be solely explained by simple CNN encoding models – rather, humans also encode them using nested combinations of discrete geometric regularities (***Sablé-Meyer et al., 2022***; ***Dehaene et al., 2022***; ***Sablé-Meyer et al., 2021***). Here, we provide direct human brain imaging evidence for the existence of two distinct cortical networks with distinct representational schemes, timing, and localization: an early occipitotemporal network well modeled by a simple CNN, and a dorso-frontal network sensitive to geometric regularity.

### Behavioral signatures of geometric regularity

At the behavioral level, the new large-scale data that we report here (n=330 participants) fully replicated our prior finding that human perception of geometric shape dissimilarity, as evaluated by a visual search task, is best predicted by a model that relies on exact geometric features, rather than by a CNN. These exact geometric feature representations can be thought of as a more abstract and compressed representations of shapes: they replace continuous variations along certain dimensions (such as angles or directions) with categorical features (right angle or not, parallel or not). Such a representation, which remains invariant over changes in size and planar rotation, turns a rich visual stimulus into a compressed internal representation.

Even in educated human adults, a small proportion of the variance in behavioral dissimilarity remained predicted by the simple CNN model, so that both CNN and geometry predictors were significant in a multiple regression of human judgments. Previously, we only found such a significant mixture of predictors in uneducated humans (whether French preschoolers or adults from the Himba community, mitigating the possible impact of explicit western education, linguistic labels, and statistics of the environment on geometric shape representation) (***Sablé-Meyer et al., 2021***). It is likely that the greater power afforded by the present experiment yielded a greater sensitivity. This finding comforts the hypothesis that two strategies for geometric shape perception are available to humans: one based on hierarchical visual processing (captured by the CNN), and the other based on an analysis of discrete geometric features – the latter dominating strongly in educated adults.

### fMRI results

Using fMRI, we found that the mere perception of geometric shapes, compared to various other visual categories, yields a reduced bilateral activation of the entire ventral visual pathway, together with a localized increased activation in the anterior IPS and in the posterior ITG. Furthermore, geometric regularity modulated fMRI activation in most of the bilateral occipito-parietal pathway, middle frontal gyrus and anterior insula. Finally, the most diagnostic result we found is that the representational similarity matrix in these regions was predicted by geometric similarity rather than by the simple CNN (Fig. 3).

The IPS areas activated by geometric shapes overlap with those active during the comprehension of elementary as well as advanced mathematical concepts (***Dehaene et al., 2022***; ***Amalric and Dehaene, 2016***, ***2019***). Although this finding must be interpreted with great caution, as activation overlap need not imply shared cognitive processes, it was confirmed at the single-subject level and agrees with the proposed involvement of these regions in an abstract, modality-independent symbolic encoding of shapes. Further support for this idea comes from the fact that these regions can be equally activated in sighted and in blind individuals when they perform mathematics (***Amalric et al., 2018***; ***Kanjlia et al., 2016***) or when they evaluate the shapes of manmade objects (***Xu et al., 2023***). Thus, the representation of shapes computed in these regions is more abstract and amodal than the hierarchy of visual filters that is thought to capture ventral visual image recognition.

One referee noted that we contrasted geometric shapes with other categories of images that may, themselves, possess some geometric features. For instance, faces possess a plane of quasisymmetry, and so do many other man-made tools and houses. Thus, our subtraction isolated the geometrical features that are present in simple regular geometric shapes (e.g. parallels, right angles, equality of length) over and above those that might already exist, in a less pure form, in other categories.

### MEG results

Using MEG, we found that even during the passive perception of simple quadrilateral shapes, in the absence of an explicit task, participants were sensitive to their geometric regularity: occasional oddball shapes elicited greater violation-of-expectation signals within blocks of regular shapes than within blocks of irregular shapes. Additionally, using RSA and source reconstruction, we observed a spatial and temporal dissociation in the processing of quadrilateral shapes: an early occipital response, well predicted by CNN models of object recognition, was followed by broadly distributed cortical activity that correlated with a model of exact geometric features. Despite the limited accuracy afforded by source reconstruction in MEG, there was considerable overlap between our RSA analysis in fMRI and in MEG.

This early response of occipital areas, followed by a later activation of a broad dorso-frontal network, may seem at odd with some results showing that, during visual image processing, the dorsal pathway may respond faster than the ventral pathway (***Ayzenberg et al., 2023***; ***Collins et al., 2019***). However, in this work, we specifically probed the processing of geometric shapes that, if our hypothesis is correct, are represented as mental expressions that combine geometrical and arithmetic features of an abstract categorical nature, for instance representing “four equal sides” or “four right angles”. It seems logical that such expressions, combining number, angle and length information, take more time to be computed than the first wave of feedforward processing within the occipito-temporal visual pathway, and therefore only activate thereafter.

### Reduced activation of the ventral visual pathway

Strikingly, our fMRI data also evidenced a hypo-activation of the ventral visual pathways to geometric shapes relative to other pictures. A large bilateral cluster of reduced activity was found when comparing shapes versus other visual categories, and a similar reduced activity was found in each of four visual areas specialized for various visual categories (faces, words, tools, and places). This reduction is compatible with our hypothesis that geometric shapes are processed differently from other pictures, and with empirical finding that a simple feedforward CNN, whose architecture usually provides a relatively good fit to inferotemporal cortex activity (***Conwell et al., 2021***; ***Kubilius et al., 2019***; ***Schrimpf et al., 2018***; ***Yamins et al., 2014***), did not provide a good model of geometric shape perception. Because our shapes consisted of a few straight lines, it is likely that they solicited the computations performed by the ventral pathway only minimally, giving rise to a lower activation when compared to more complex visual stimuli such as pictures of faces or houses.

### Two Visual Pathways

Brain-imaging studies of the ventral visual pathway indicate that it is subdivided into patches responding to different visual categories such as faces, tools or houses, with a partial correspondence between human and non-human primates (***Arcaro et al., 2017***; ***Margalit et al., 2020***; ***Rauschecker et al., 2012***; ***Tsao and Livingstone, 2008***; ***Khosla et al., 2022***; ***Allen et al., 2021***; ***Kriegeskorte et al., 2008b***). Converging evidence from behavior (***Biederman and Ju, 1988***) and neuroimaging (***Ayzenberg et al., 2022b***; ***Lescroart and Biederman, 2013***; ***Papale et al., 2019***, ***2020***) has underlined the central role of the shape of objects, contour and texture in object recognition. A basic complementarity has been identified between the ventral visual pathway, crucial for visual identification, and the dorsal occipito-parietal route, extracting visuo-spatial information about orientation and motor affordances. However, this simplified view needs to be nuanced by evidence of shape processing in the dorsal pathway as well, especially in posterior areas (***Freud et al., 2016***; ***Xu, 2018***), in both human and non-human primates. More specifically, recent work challenges the idea that the shape of objects is computed solely in the ventral pathway, and instead suggests that the dorsal pathway, and in particular the IPS, may first compute global shape information and then propagate it to the ventral pathway for further processing (***Ayzenberg and Behrmann, 2022***). Our results are compatible with this new look at dorsal/ventral interactions: in our fMRI localizer, geometric shapes elicited reduced ventral and increased dorsal activation when compared to other visual categories. Geometric shapes could be considered as conveying very pure information about global shape, entirely driven by contour geometry, and fully devoid of other information such as texture, shading, or curvature. More research is needed to understand the dynamic collaboration between the ventral and dorsal streams during shape recognition, but the present results, as well as the arguments put forward in (***Ayzenberg and Behrmann, 2022***) concur to suggest that shape identification does not solely rely on the ventral visual pathway.

Interestingly, recent work shows that within the visual cortex, the strongest relative difference in cortical surface expansion between human and non-human primates is localized in parietal areas (***Meyer et al., 2025***). If this expansion reflected the acquisition of new processing abilities in these regions, it might explain the observed differences in geometric abilities between human and non-human primates (***Sablé-Meyer et al., 2021***).

### Lateral Occipital Cortex

Of special relevance to this work is the role of the Lateral Occipital Cortex (LOC) in shape perception (***Grill-Spector et al., 2001***). The posterior ITG activation that we observed lies just anterior to the LOC and possibly overlapped with it. The LOC has been repeatedly associated with shape processing (***Grill-Spector et al., 1998***; ***Kanwisher et al., 1996***; ***Kourtzi and Kanwisher, 2000***; ***Malach et al., 1995***), and our work converges to suggest that this region and the cortex just anterior to it play a key role in the perception of simple shapes in a way that is invariant to 2D and 3D rotations (***Kourtzi et al., 2003***). However, unlike ours, previous work focused primarily on irregular potatoe-like shapes or objects and their contours.

Object selectivity in the LOC is already present in early infancy, with a sensitivity to shape and not texture already observed at 6-months of age (***Emberson et al., 2017***). At age 5-10, size invariance, but not viewpoint invariance, has been established (***Nishimura et al., 2015***). However, to our knowledge, no study has targeted geometrically regular shapes, and in particular how changes in regularity impacts LOC activity. While some of our quadrilateral stimuli can be constructed as 3D projections of one another (either by perspective or by orthogonal projection), this should make them more similar to each other within the LOC given its object viewpoint invariance. Further work should use within-subject LOC localizers such as objects minus scrambled-objects to establish the exact cortical relation between the present geometric regularity effect and the classical LOC region. We speculate that the more anterior part of lateral occipito-temporal cortex may extract more abstract geometric descriptors of shapes than the classical LOC, as also suggested by its sensitivity to mathematical training (***Amalric and Dehaene, 2016***).

### Development of geometry representations

Interestingly, the IPS and posterior ITG activations to geometric shapes were consistently observed at very similar locations in adult and 6-year-olds. Furthermore, the overlap with a math-responsive network was present in both age groups, with ROIs that respond to number more than words also responding to geometric shapes more than other visual categories. This finding fits with previous evidence of an early responsivity of the IPS to numbers and arithmetic in young children prior to formal schooling (***Ansari, 2008***; ***Cantlon and Li, 2013***; ***Izard et al., 2008***). In agreement with the existence of a behavioral geometric regularity effect in preschoolers (***Sablé-Meyer et al., 2021***), we hypothesize that an intuitive encoding of number and geometric features precede schooling and possibly guide the subsequent acquisition of formal mathematics (***Izard et al., 2022***).

On the other hand, the fMRI results from the intruder task were much less stable in children, and neither the geometric regularity effect nor the geometry-based RSA analysis yielded strong results at the whole-brain level (only one ventromedial cluster was associated with the regularity effect, see Figure S3). In previous work, we have shown that besides an overall decrease in performance between adults and first graders, the profile of behavior across different shapes is partially distinct in preschoolers: in them, both the CNN and the symbolic model are on par in predicting behavior, suggesting that unlike adults, children frequently mix the two strategies for identifying geometric shapes. Such a mixture of strategies could explain why the correlation with our empirical estimation of complexity, based on adult data, yielded a weaker and more noisy correlation in children. The RSA analysis is compatible with this: in the group analysis, while the CNN predictor was significant in the visual cortex of children, the symbolic geometry model did not reach significance anywhere. Future research with a larger number of participants could attempt to sort out children as a function of whether their behavior is dominated by geometric features or by the CNN model, and then examine how their brain activity profiles differ.

### Artificial Neural Networks: limits and promising directions

While simple CNNs are predictive of early ventral visual activity, the present work adds to a growing list of their limits as full models of visual perception (***Bowers et al., 2022***; ***Jacob et al., 2021***). Simple CNNs have been shown to fail to model many visual illusions (***Bowers et al., 2022***; ***Jacob et al., 2021***), geometrically impossible objects (***Heinke et al., 2021***), global shape, part-whole relations and other Gestalt properties (***Bowers et al., 2022***). While larger CNNs may overcome some of these issues, and indeed here we found that the ConvNeXT CNN-like architecture could mimic our MEG data, their huge training sets likely include human geometric shapes and drawings, without which they may fail in out-of-domain generalization (***Mayilvahanan et al., 2025***). Human children, by contrast, recognize and produce geometric drawings early on and with little or no explicit training (***Goodenough, 1928***; ***Long et al., 2024***; ***Saito et al., 2014***). Thus, these observations suggest that, to capture the human sense of geometry, current artificial-intelligence models may have to be supplemented, not only by increasing their size (***Conwell et al., 2021***) and training sets, but also by changing their architecture in ways that better incorporate findings from psychology and neuroscience (***Thompson et al., 2023***; ***Biederman, 1987***). Indeed, connectionist models based on the transformer architecture seem to be superior in capturing human geometric perception (***Campbell et al., 2024***), and indeed we found that our results could also be captured by the late layers of visual transformer DINOv2 (***Oquab et al., 2023***)see Fig. S5. and Fig. S6). This finding may offer an exciting avenue to analyze a mechanistic implementation of symbol-like representations in a connectionist architecture. However, it is also possible that much more sophisticated mechanisms of Bayesian program inference are required to truly capture the remarkable efficiency with which humans grasp abstract shapes and geometric drawings (***Lake et al., 2015***, ***2016***; ***Sablé-Meyer et al., 2022***).

### Shape Skeleton and Principal Axis

Skeletal representations have been proposed as human-like representations of visual shapes, particularly appropriate for biological shapes such as animals or plants (***Blum, 1973***). Even for simple geometric shapes, skeletal representation may be automatically and unconsciously computed (***Firestone and Scholl, 2014***). However, in the present work, the two skeletal shape representations we tested did not model human data well (see Fig. S5 and Fig. S6). Within our quadrilaterals, which are visually similar to one another, all but the square possess the same skeleton topological graph, with only variations in the length and angles of the skeletal segments. Thus, a skeletal representation is probably not the source of the large geometric regularity effect that we observed here and in past work (***Sablé-Meyer et al., 2021***). Still, as previously argued (Sablé-Meyer et al., 2022), geometric and medial axis theories need not be seen as incompatible. Rather, we conceive of them as complementary codes, best suited for distinct shape domains. Even after extraction of a figure’s skeleton, further compression could be achieved by encoding its geometric regularity, and this may be what humans do when they draw a snake, for instance, as a geometrically regular zigzag.

### Neurophysiological implementation of geometry

Symbolic models are often criticized for lack of a plausible neurophysiological implementation. It is therefore important to discuss whether and how the postulated symbolic geometric code could be realized in neural circuits. There are several distinct challenges. First, some neurons should encode individual features such as lines and curves, or features formed by their relationships (e.g. parallelism, right-angle). Second, these codes should be discrete and categorical, forming sharp boundaries between, say, parallel versus non-parallel lines. Third, they should enter into compositional expressions describing how individual features combine into the whole shape (e.g. “a shape with four equal sides and four equal right-angles”).

The first point has been studied in both humans and monkeys in the context of research on the non-accidental properties (NAPs) of objects. NAPs are qualitative features of object shapes, such as straight versus curved, which remain invariant when an object is rotated. Metric properties (MPs), on the other hand, are quantitative properties such as amount of curvature that do not exhibit such invariance. Behavioral research has demonstrated that, for equivalent amounts of pixel change in the image, changes in NAPs are more discriminable than changes in MPs, in educated and uneducated human adults, toddlers and even in infants (***Amir et al., 2012***; ***Biederman et al., 2009***; ***Kayaert and Wagemans, 2010***). Furthermore, the firing of neurons in monkey infero-temporal cortex is more sensitive to changes in NAPs than in MPs, whether they are conveyed by 3D shapes (***Kayaert et al., 2003***) or 2D shapes (***Kayaert et al., 2005a,b***). These findings occurred even in the absence of a task other than passive fixation, and without particular training for the stimuli. Further work with 2D shapes, including triangles and quadrilaterals partially overlapping with the present stimuli, showed that IT cortex neurons could also be sensitive to more global shape properties such as axis curvature or “taper” (the difference between a rectangle seen upfront or at a slanted axis) (***Kayaert et al., 2005a***). Multidimensional scaling of macaque neural population responses organized the tested shapes in a systematic “shape space” where, for instance, the rectangle occupies a extreme corner, thus making it distinct from its curved or tapered variants (***Kayaert et al., 2005a***, figure 5).

While these previous non-human primate findings may provide a neurophysiological basis for the fMRI and MEG responses observed here, the neural implementation of our third requirement of having neural codes enter compositional expressions remains elusive. What is more, there are still notable differences between humans and macaques: humans do not need to be extensively trained to understand the differences between geometric shapes (***Izard et al., 2022***; ***Izard and Spelke, 2009***), and spontaneously impose geometric categories such as “parallel” or “right angle” to a continuum of angles between lines (***Dillon et al., 2019***). Several behavioral findings suggest that a distinct neural code for geometry may exist in humans. Non-human primates perform poorly on a broad variety of perceptual and production tasks with geometric shapes: baboons do not exhibit a human-like geometric complexity effect (***Sablé-Meyer et al., 2021***; ***Dehaene et al., 2022***); chimpanzees do not transfer learning of visual categories from concrete pictures to geometric line drawings (***Close and Call, 2015***); and chimpanzees behave very differently from children in free-drawing experiments where they have to complete partial line-drawings of faces (***Saito et al., 2014***). These findings suggests that an understanding of the mechanisms that underlie the human coding of geometric shapes may ultimately shed light on the cognitive and neural singularity of the human brain (***Dehaene et al., 2022***).

In the future, replicating the present experiments with monkey fMRI and electrophysiology is therefore an important goal. The methodology could also be extended to the perception of a broader set of geometric patterns (circles, spirals, crosses, zigzags, plaids, etc) which recur since prehistory and in the drawings of children and adults of various cultures, thus testing whether they too originate in a minimal and universal “language of thought” (***Sablé-Meyer et al., 2022***) and whether such a language is unique to the human species (***Dehaene, 2026***; ***Dehaene et al., 2022***). Finally, this research should ultimately be extended to the representation of 3-dimensional geometric shapes, for which similar symbolic generative models have been proposed (***Biederman, 1987***; ***Leyton, 2003***).

## Method

### Experiment 1

#### Participants

330 participants (142 Females, 177 Males, 11 Others; mean age 51.1 years, SD=16.8) were recruited via a link provided on a New York Time article (available at https://www.nytimes.com/2022/03/22/science/geometry-math-brain-primates.html) which reported previous research from the lab and featured a link to our new online experiment. When participants clicked the link, they landed on a page with our usual procedure for online experiments, including informed consent and demographic questions. No personal identity information was collected.

#### Task

On each trial, participants were shown a 3×3 square grid of shapes, eight of which were copies of the same shape up to rotation and scaling, and one of which was a different shape. Participants were asked to detect the intruder shape by clicking on it. Auditory feedback was provided in the form of tones of ascending or descending pitch, as well as coloring of the shapes (the intruder was always colored in green indicating what the right answer was, and in case of an erroneous choice, the chosen shape was colored in red indicating a wrong answer).

#### Stimuli

Shapes design followed previous work exactly (***Sablé-Meyer et al., 2021***): eleven quadrilaterals with a varying number of geometric features, matched for average pairwise distance of all vertices and length of the bottom side. Shapes were presented in pure screen white on pure screen black. Each shape was differently scaled and rotated by sampling nine values without replacement in the following scaling factors [0.85, 0.88, 0.92, 0.95, 0.98, 1.02, 1.05, 1.08, 1.12, 1.15] and rotation angles [-25°, −19.4°, −13.8°, −8.3°, −2.7°, 2.7°, 8.3°, 13.8°, 19.4°, 25°]. Note that the rotation angles were centered on 0° but excluded this value so that the sides of the shapes were never strictly vertical or horizontal. Participants performed 110 trials (11×10), one for each pair of different reference and intruder shapes. The order of trials was randomized, subject to the constraint that no two identical reference shapes were used on consecutive trials, and that the outlier of a trial was always different from the reference shape of the previous trial. Two examples of trials are shown in Fig.1B.

#### Procedure

The experimental procedure started with instructions, followed by a series of questions: device used (mouse or touchscreen), country of origin, gender, age, highest degree obtained. Participants then provided subjective self-evaluation assessments, with answers on a Likert scale from 1 to 10, for the following items: current skills in mathematics; current skills in first language. Finally, participants performed the task. The instructions text was the following: “The game is very simple: you will see sets of shapes on your screen. Apart from small rotation and scaling differences, they will be identical, except for one intruder. Your task is to respond as fast and accurately as you can about the location of the intruder by clicking on it. The difficulty will vary, but you always have to answer.”

#### Estimation of empirical representational dissimilarity

To estimate the representational dissimilarity across shapes, first we estimated the dissimilarity between two shapes as the average success rate divided by the average response time. Indeed, if two shapes are very dissimilar, we expect participants to make few mistakes (high success rate) and find the intruder fast (low response time), yielding a high value, and vice versa. Because we didn’t have predictions using the asymmetry in visual search (e.g. finding a square within rectangles versus a rectangle versus squares), we then averaged over these paired conditions, thereby turning the square dissimilarity matrix into a triangular dissimilarity matrix. Finally, we z-scored these dissimilarity estimates.

As participants performed a single trial per pair of shapes, the estimation of the dissimilarity is noisy at the single participant level (either 0 or 1/RT depending on whether they answered correctly). We kept this estimate for analyses at the single participant level or mixed-effect analysis; however, for analyses that required a single RDM estimate across participants, we pooled the data from participants to estimate the average success rates and response times, hence before estimating the empirical RDMS, rather than estimate one RDM per participant and then averaging.

#### Comparison with Model RDMs

To obtain theoretical representational dissimilarity matrices (RDMs) with which to compare the present behavioral data, as well as subsequent brain-imaging data, we proceeded as follows. For the CNN encoding model, we downloaded from GitHub the weights for several neural networks (CORnet (***Kubilius et al., 2018***), ResNet (***He et al., 2016***) and DenseNet (***Huang et al., 2018***); all high scoring on brain-score (***Schrimpf et al., 2018***)), all pre-trained with ImageNet and not specifically trained for our task. We extracted the activation vectors in each hidden layer associated to each shape with the 6×6=36 different orientations and scaling used in the experiment. For each shape, we separated their 36 exemplars randomly into two group to have independent estimation of the representation vectors from the network, and used the cross-validated Mahalanobis distance between these two splits to estimate the distance between each pair of shapes. This provides us with an RDM that captures how dissimilar shapes are according to their internal representations in a CNN of object recognition. Unless specified otherwise (see supplementary text), we report correlation with CORnet layer IT following (***Sablé-Meyer et al., 2021***).

For the geometric feature model, we estimate a feature vector of geometric features for each shape. This feature vector includes information about (i) right angles (one for each angle, 4 features); (ii) angle equality (one for each pair of angles, 6 features); (iii) side length equality (one for each pair of sides, 6 features); and (iv) side parallelism (one for each pair of sides, 6 features); all of this was done up to a tolerance level (e.g. an angle slightly off a right angle still counts as a right angle), using the tolerance value of 12,5% which was fitted previously to independent behavioral data (***Sablé-Meyer et al., 2021***). The dissimilarity between each pair of shapes was the difference between the number of features that each shape possesses: because both the square and the rectangle share many features, they are similar. Two very irregular shapes also end up similar as well. Conversely, the square and an irregular shape end up very dissimilar.

### Experiment 2

#### Participants

Twenty healthy French adults (9 females; 19-37 years old, mean age = 24.6 years old, SD: 5.2 years old) and 25 French first graders (13 females; all 6 years old) participated in the study. Three children quit the experiment before any task began because they did not like the MRI noise or lying in the confined space for the scanner. One child completed the localizer task but not the other tasks. One child was missing a single run from the intruder task. All participants had normal hearing, normal or corrected-to-normal vision, and no known neurological deficit. All adults and guardians of children provided informed consent, and adult participants were compensated for their participation.

#### Task

In three localizer runs, children and adult participants were exposed to eight different image categories, such that single geometric shapes could be compared to matched displays of faces, houses and tools, and rows of three geometric shapes could be compared to matched displays of numbers, French words, and Chinese characters. To maintain attention, participants were asked to keep fixating on a green central fixation dot (radius=8pixels, RGB color=26, 167, 19, always shown on the screen), and to press a button with their right-hand whenever a star symbol was presented. The star spanned roughly the same visual angle as the stimuli from the eight categories, and appeared randomly once in one of the two blocks per category (8 target stars total), between the 3rd to the 6th stimuli within that block. As feedback, a 300 ms 650 Hz beep sound was provided after each button press.

#### Stimuli

In each miniblock, participants saw a series of 6 grayscale images, one per second, belonging to one of eight different categories: faces, houses, tools, numbers, French words, Chinese characters, single geometric shapes, and rows of three geometric shapes. Each category comprised 20 exemplars. All faces, 16 houses, and 18 tools had been used in previous localizer experiments (***Zhan et al., 2018***). For face stimuli, front-view neutral faces (20 identities, 10 males) were selected from the Karolinska Directed Emotional Faces database (***Lundqvist et al., 1998***). The stimuli were aligned by the eyes and the iris distances. A circular mask was applied to exclude the hair and clothing below the neck. House and tool stimuli were royalty-free images obtained from the internet. House stimuli were photos of 2 to 3-story residence houses. Tool stimuli were photos of daily hand-held tools: half of the images were horizontally flipped, so that there were 10 images in a position graspable for the left and right hand respectively. For French word stimuli, 3-letter French words were selected which were known to first graders and had high occurrence frequencies (range=7-2146 occurrences per million, mean=302, SD=505, based on Lexique, http://www.lexique.org/). Chinese characters were selected from the school textbook of Chinese first graders. Chinese word frequency (range=11-1945 occurrences per million, mean=326, SD=451 (***Cai and Brysbaert, 2010***)) was not significantly different from French words used here (t(38)=.2, p=.87). Single-digit formula stimuli were 3-character simple operations in the form of “x+y” or “x-y” with x greater than y, x ranging from 2 to 5, and y from 1 to 4. Single shapes consisted in a single, centered outline of a geometrical shape (diamond, hexagon, rhombus, parallelogram, rectangle, square, trapezoid, isosceles triangle, equilateral triangle and right triangle), and were matched in luminance, contrast, and visual angle to the faces/houses/tools/words/Chinese characters which also displayed single objects. A row of shapes consisted of three different shapes side by side, whose total width, size, and line width were matched with 3-letter French words and 3-character single-digit operations. To match the appearance of the monospaced font in previous work (***Vinckier et al., 2007***), the monospaced font Consolas was used for the French words and numbers, with identical font weight 900. The font for Chinese characters was Heiti, which looks similar to Consolas. Random font size (uniform in 35-55px font size) were repeatedly sampled until text stimuli achieved similar variability as with the other categories.

The stimuli were embedded in a gray circle (RGB color=157, 157, 157, radius=155 pixels), on the screen with a black background. Within the gray circle, the mean luminance and contrast of the 8 stimuli categories did not differ significantly (luminance: F(7,152)=.6, p=.749; contrast: F(7,152)=1.2, p=.317), see Fig.S2

#### Procedure

The eight categories were presented in distinct blocks of 6s each, fully randomized for block presentation order, with the restriction that there were no consecutive blocks from the same category. Each miniblock comprised 6 stimuli, in random order. Each stimulus was presented for 1s, with no interval in between (***Dehaene-Lambertz et al., 2018***). The inter-block interval duration was jittered (4, 6, or 8s; mean = 6s). Each of the eight-block types appeared twice within each run. A 6s fixation period was included at both the beginning and the end of the run. Each run lasted for 3min 24s, and participants performed three such runs during the fMRI session.

### Experiment 3

#### Participants

Identical to experiment 2

#### Procedure and Stimuli Task

We adapted the geometric intruder detection task (***Sablé-Meyer et al., 2021***; ***Dehaene et al., 2006a***) to the fMRI scanner. On each trial, participants (children and adults) saw an array of 6 shapes around fixation (3 on the right, and 3 on the left; see Fig.3A). Five shapes were identical except for a small amount of random rotation and scaling, while one was a deviant shape. Because the pointing task used in (***Sablé-Meyer et al., 2021***) was not possible in the limited space of the fMRI scanner, participants were merely asked to click a button with their left or right hand, thereby indicating on which side they thought the deviant was. Participants responded on every trial, the side of the correct response was counterbalanced within each shape, and we verified that the average motor response side was unconfounded with geometric shape or complexity. After each answer, auditory feedback was provided with a tone of high, increasing pitch when the answer was correct, and a low-pitch tone otherwise.

#### Stimuli

Geometric shapes were generated following the procedure described in previous work (***Sablé-Meyer et al., 2021***): to fit the experiment within the time constraints of children fMRI, a subset of shapes was used, comprising square, rectangle, isosceles trapezoid, rhombus, right hinge and irregular shapes to span the range of complexity found in (***Sablé-Meyer et al., 2021***). Following previous work (***Sablé-Meyer et al., 2021***), deviants were generated by displacing the bottom right corner by a constant distance in four possible positions (see Fig.3A). That distance was a fraction of the average distance between all pairs of points, which was standardized across shapes (45% change). On each trial, six gray-on-black shapes were shown (shape color rgb values: 127,127,127). Shapes were displayed along two semicircles: the positions were determined by positioning the three leftmost (resp. rightmost) shapes on the left side (resp. right side) of a circle of radius 120px, at angles 0, pi/2 and pi, and then shifting them 100px to the left (resp. right). The rotation and scaling of each shape were randomized so that no two shapes had the same scaling or rotation factor, and values were sampled in [0.875, 0.925, 0.975, 1.025, 1.075, 1.125] for scaling and [-25°, −15°, −5°, 5°, 15°, 25°] for rotations, avoiding 0° to prevent alignment of specific shapes with screen borders. One of the shapes was an outlier, whose position was sampled uniformly in all six possible positions such that no two consecutive trials featured outliers in the same position. Outliers were sampled uniformly from the four possible types of outliers, so that all outlier types occurred as often, but no two consecutive trials featured identical outlier types.

#### Procedure

The six shapes were presented in miniblocks, in randomized order, with no two consecutive blocks with the same type of shape. Each block comprised 5 consecutive trials with an identical base shape, each with 2s of stimulus presentation and 2s of fixation. There was a 4s, 6s or 8s delay between blocks. A central green fixation cross was always on display, and it turned bold 600 ms before a block would start. Each run of the outlier detection task lasted 3m40s.

### fMRI methods common to experiments 2 and 3

#### MRI Acquisition Parameters

MRI acquisition was performed on a 3T scanner (Siemens, Tim Trio), equipped with a 64-channel head coil. Exactly 113 functional scans covering the whole brain were acquired for each localizer run, as well as on 179 functional scans covering the whole brain for each run of the geometry task. All functional scans were using a T2*-weighted gradient echo-planar imaging sequence (69 interleaved slices, TR = 1.81 s, TE = 30.4 ms, voxel size = 2×2×2mm, multiband factor = 3, flip angle = 71 degrees, phase encoding direction: posterior to anterior). To reconstruct accurate anatomical details, a 3D T1-weighted structural image was also acquired (TR = 2.30 s, TE = 2.98 ms, voxel size = 1×1×1mm, flip angle = 9 degrees). To estimate distortions, two spin-echo field maps with opposite phase encoding directions were acquired: one volume in the anterior-to-posterior direction (AP) and one volume in the other direction (PA). Each fMRI session lasted for around 50min for children including (in order) 3 runs of a task not discussed here, 3 Category localizer runs, T1 collection, and 2 Geometry runs. For adults, the session lasted for around 1h 20min because they took the same runs as for children as well as an additional harder version of the geometry task. This version, which involved smaller deviant distances and a stimulus presentation duration of only 200 ms, turned out to be too difficult. While the overall performance still shows a correlation with complexity (r²=.63, p<.02), it was entirely driven by two shapes, the square and the rectangle: other shapes were equally hard, and although they were better than chance they did not correlate with complexity (r^2^=.35,p=.22) while the correlations remained for the simpler condition.

#### Data analysis

Preprocessing was performed with the standard pipeline fMRIPrep. Results included in this manuscript come from preprocessing performed using fMRIPrep 20.0.5 (***Esteban et al., 2018a,b***), which is based on Nipype 1.4.2 (***Gorgolewski et al., 2011***, ***2018***), and generated the following detailed method description.

#### Anatomical Data

The T1-weighted (T1w) image was corrected for intensity non-uniformity (INU) with N4BiasFieldCorrection (***Tustison et al., 2010***), distributed with ANTs 2.2.0 (***Avants et al., 2008***), and used as T1w-reference throughout the workflow. The T1w-reference was then skull-stripped with a Nipype implementation of the antsBrainExtraction.sh workflow (from ANTs), using OASIS30ANTs as target template. Brain tissue segmentation of cerebrospinal fluid (CSF), white-matter (WM) and gray-matter (GM) was performed on the brain-extracted T1w using fast (***Zhang et al., 2001***). Brain surfaces were reconstructed using recon-all (***Dale et al., 1999***), and the brain mask estimated previously was refined with a custom variation of the method to reconcile ANTs-derived and FreeSurfer-derived segmentations of the cortical gray-matter of Mindboggle (***Klein et al., 2017***). Volume-based spatial normalization to two standard spaces (MNI152NLin6Asym, MNI152Nlin2009cAsym) was performed through nonlinear registration with antsRegistration (ANTs 2.2.0), using brain-extracted versions of both T1w reference and the T1w template. the following templates were selected for spatial normalization: FSL’s MNI ICBM 152 non-linear 6^th^ Generation Asymmetric Average Brain Stereotaxic Registration Model (***Evans et al., 2012***) and ICBM 152 Nonlinear Asymmetrical template version 2009c (***Fonov et al., 2009***).

#### Functional Data

For each of the 10 BOLD EPI runs found per subject (across all tasks and sessions), the following preprocessing was performed. First, a reference volume and its skull-stripped version were generated using a custom methodology of fMRIPrep. Susceptibility distortion correction (SDC) was omitted. The BOLD reference was then co-registered to the T1w reference using bbregister (FreeSurfer) which implements boundary-based registration (***Greve and Fischl, 2009***). Co-registration was configured with six degrees of freedom. Head-motion parameters with respect to the BOLD reference (transformation matrices, and six corresponding rotation and translation parameters) are estimated before any spatiotemporal filtering using mcflirt (***Jenkinson et al., 2002***). BOLD runs were slice-time corrected using 3dTshift from AFNI 20160207 (***Cox and Hyde, 1997***). The BOLD time-series (including slice-timing correction when applied) were resampled onto their original, native space by applying the transforms to correct for head-motion. These resampled BOLD time-series will be referred to as preprocessed BOLD in original space, or just preprocessed BOLD. The BOLD time-series were resampled into several standard spaces, correspondingly generating the following spatially-normalized, preprocessed BOLD runs: MNI152Nlin6Asym, MNI152Nlin2009cAsym. First, a reference volume and its skull-stripped version were generated using a custom methodology of fMRIPrep. Several confounding time-series were calculated based on the preprocessed BOLD: framewise displacement (FD), DVARS and three region-wise global signals. FD and DVARS are calculated for each functional run, both using their implementations in Nipype (following the definitions by (***Power et al., 2014***)). The three global signals are extracted within the CSF, the WM, and the whole-brain masks. Additionally, a set of physiological regressors were extracted to allow for component-based noise correction (***Behzadi et al., 2007***). Principal components are estimated after high-pass filtering the preprocessed BOLD time-series (using a discrete cosine filter with 128s cut-off) for the two CompCor variants: temporal (tCompCor) and anatomical (aCompCor). tCompCor components are then calculated from the top 5% variable voxels within a mask covering the subcortical regions. This subcortical mask was obtained by heavily eroding the brain mask, which ensures it does not include cortical GM regions. For aCompCor, components are calculated within the intersection of the aforementioned mask and the union of CSF and WM masks calculated in T1w space, after their projection to the native space of each functional run (using the inverse BOLD-to-T1w transformation). Components are also calculated separately within the WM and CSF masks. For each CompCor decomposition, the k components with the largest singular values are retained, such that the retained components’ time series are sufficient to explain 50 percent of variance across the nuisance mask (CSF, WM, combined, or temporal). The remaining components are dropped from consideration. The head-motion estimates calculated in the correction step were also placed within the corresponding confounds file. The confound time series derived from head motion estimates and global signals were expanded with the inclusion of temporal derivatives and quadratic terms for each(***Satterthwaite et al., 2013***). Frames that exceeded a threshold of 0.5 mm FD or 1.5 standardised DVARS were annotated as motion outliers. All resamplings can be performed with a single interpolation step by composing all the pertinent transformations (i.e. head-motion transform matrices, susceptibility distortion correction when available, and co-registrations to anatomical and output spaces). Gridded (volumetric) resamplings were performed using antsApplyTransforms (ANTs), configured with Lanczos interpolation to minimize the smoothing effects of other kernels(***Lanczos, 1964***). Non-gridded (surface) resamplings were performed using mri_vol2surf (FreeSurfer).

Many internal operations of fMRIPrep use Nilearn 0.6.2 (***Abraham et al., 2014***), mostly within the functional processing workflow.

### fMRI GLM Models

fMRI first-level models (general linear models or GLM) were computed by convolving the experimental design matrix (specific to each task, see below) with SPM’s HRF model as implemented in Nilearn, with the following parameters: spatial smoothing using a full width at half maximum window (fwhm) of 4mm; a second-order autoregressive noise model; and signal standardized to percent signal change relative to whole-brain mean. The following confound regressors were added: polynomial drift models from constant to 5^th^ order (6 regressors); estimated head translation and rotation on three axes (6 regressors) as well as the following confound regressors given by fmriprep: average cerebro-spinal fluid signal (1 regressor), average white matter signal (1 regressor), and the first five high-variance confounds estimates (***Behzadi et al., 2007***) (5 regressors).

In the visual category localizer, the design matrix contained distinct events for each visual stimulus, grouped by category, each with a duration of 1s, including a specific one for the target star, leading to 9 regressors (Chinese, face, house, tools, numbers, words, single shape, three shapes, and star). In the geometry task runs, each reference shape was associated to a regressor with trial duration set to 2s, thus leading to 6 regressors (square, rectangle, rhombus, iso-trapezoid, hinge and random). In both cases, button presses were not modeled as they were fully correlated with predictors of interest (either the star for the localizer, or every single trial for the geometry task).

Second-level models were estimated after additional smoothing with a full width at half maximum window of 8mm. Statistical significance of clusters was estimated with a bootstrap procedure as follows: given an uncorrected p-value of .001, clusters were identified by contiguity of voxels that had p-values below this threshold. Then, the p-value of a cluster was derived by comparing its statistical mass (the sum of its t-values) to the distribution of the maximum statistical mass obtained by performing the same contrast after randomly swapping the sign of each participant’s statistical map. We performed this swapping 10,000 times for each contrast we estimated, and computed the corrected p-value accordingly; for instance, a cluster whose mass was only outperformed by 3 random swaps out of the 10,000 was assigned a p-value of .0003.

#### Searchlight RSA analyses

First, we estimated the Representational Dissimilarity Matrix (RDM) within spheres centered on each voxel. For this we performed a searchlight sweep across the whole brain. We extracted the GLM coefficients of the geometric shapes from each voxel and all the neighboring voxels in a 3-voxel radius (=6mm), for a total of 93 voxels per sphere. We discarded voxels where more than 50% of this sphere fell outside the participant’s brain. We extracted the betas of each shape and used a cross-validated Mahalanobis distance across runs (crossnobis, implemented in rsatoolbox) to estimate the dissimilarity between each pair of shapes. We attributed this distance to the center of the searchlight, thereby estimating an empirical RDM at each location.

Then we compared this empirical RDM with the two RDMs derived from our two models separately, using a whitened correlation metric. Both choices of metrics (crossnobis and whitened correlation) follow the recommendations from a previous methodological publication (***Diedrichsen et al., 2021***).

Finally, we computed group-level statistics by smoothing the resulting correlation map (fwhm=8mm) and performing statistical maps, cluster identification, and statistical inference at the cluster level as we did for the second-order level analysis, but with a one-tailed comparison only as we did not consider negative correlations.

The bar plot for ROIs in figure S4 reflect a subject-specific voxel localization within ROIs. Within each ROI identified we find, for each subject, the 10% most responsive subject-specific voxels in the same contrast used to identify the cluster. To avoid double-dipping, we selected these voxels using the contrast from one run, then collected the fMRI responses (beta coefficients) from the other runs; we perform this procedure across all runs and average the responses. Error bars indicate the standard error of the mean across participants.

### Experiment 4

#### Participants

Twenty healthy French adults (13 females; 21-42 years old, mean: 24.9 years old, SD: 8.1 years old) participated in the MEG study. All participants had normal hearing, normal or corrected-to-normal vision, and no neurological deficit. All adults provided informed consent and were compensated for their participation. For all but one participant, we had access to anatomical recordings in 3T MRI, either from prior, unrelated experiments in the lab, or because the MEG session was immediately followed by a recording. Because of one participant missing an anatomical recording, analyses that required source reconstruction were performed on nineteen subjects.

#### Task

During MEG, adult participants were merely exposed to shapes while maintaining fixation and attention. As in previous work (e.g. ***Al Roumi et al., 2023***; ***Benjamin et al., 2024***), the goal was to examine the spontaneous encoding of stimuli and the presence or absence of a novelty response to occasional deviants.

#### Stimuli

All 11 geometric shapes in Fig.1A were presented in miniblocks of 30 shapes. Geometric shapes were presented centered on the screen, one shape every second, with shapes remaining onscreen for 800ms and a centered fixation cross present between shapes for 200ms. To make the shapes more attractive (and since the same shape were also used in an infant experiment, not reported here), during their 800ms presentation, the shapes slowly increased in size: in total, a scale factor of 1.2 was applied over the course of 800ms, with linear interpolation of the shape size during the duration of the presentation. Shapes were presented in miniblocks following an oddball paradigm: within a miniblock, all shapes were identical up to scaling (randomly sampled in [0.875, 0.925, 0.975, 1.025, 1.075, 1.125]) and rotation (sampled in [-25°, −15°, −5°, 5°, 15°, 25°]), except for occasional oddballs which were deviant version of the reference shape. Each miniblock comprised 30 shapes, 4 of which were oddballs that could replace any shape after the first six. Two oddballs never appeared in a row. There was no specific interval between miniblocks beyond the usual duration between shapes. A run was made of 11 miniblocks, one per shape in random order, and participants attended 8 runs except two participants who are missing a single run due to experimenters’ mistakes when setting up the MEG acquisition.

#### Procedure

After inclusion by the lab’s recruiting team, participants were prepared for the MEG with electrocardiogram (ECG) and electrooculogram (EOG) sensors, as well as four Head Position Indicator (HPI) coils, which were digitalized to track the head position throughout the experiment. Then we explained participants that the task was a replication of an experiment with infants, and therefore was purely passive: they would be shown shapes and were instructed to pay attention to each shape, while trying to avoid blinks as well as body and eye movements. They sat in the MEG and we checked the head position, ECG/EOG and MEG signal. From that point onward, we never opened the MEG door again to avoid resetting MEG signals and allow for optimal cross-run decoding and generalization. Participants took typically 8 runs consecutively, with small breaks with no stimuli between runs to rest their eyes. At the end of the experiment, participants took the intruder test from previous work(***Sablé-Meyer et al., 2021***) on a laptop computer outside of the MEG, and finally we spent some time debriefing with participants the goal of the experiment.

To ensure high accuracy of the timing in the MEG, each trial’s first frame contained a white square on the bottom of the screen, which was hidden from participants but recorded with a photodiode. The same area was black during the rest of the experiment. The ramping up of the photodiode was therefore synchronized with the screen update and the appearance of the stimulus, ensuring robust timing for analyses. Then each “screen update” event was linked to a recorded list of presented shapes.

#### MEG Acquisition Parameters

Participants were instructed to look at a screen while sitting inside an electromagnetically shielded room. The magnetic component of their brain activity was recorded with a 306-channel, whole-head MEG by Elekta Neuromag© (Helsinki, Finland). The MEG helmet is composed of 102 triplets of sensors, each comprising one magnetometer and two orthogonal planar gradiometers. The brain signals were acquired at a sampling rate of 1000 Hz with a hardware high-pass filter at 0.03 Hz.

Eye movements and heartbeats were monitored with vertical and horizontal electro-oculograms (EOGs) and electrocardiograms (ECGs). Head shape was digitized using various points on the scalp as well as the nasion, left and right pre-auricular points (FASTTRACK, Polhemus). Subjects’ head position inside the helmet was measured at the beginning of each run with an isotrack Polhemus Inc system from the location of four coils placed over frontal and mastoidian skull areas.

#### Preprocessing of MEG signals

The preprocessing of the data was performed using `MNE-BIDS-Pipeline`, a streamlined implementation of the core ideas presented in the literature (***Jas et al., 2018***) and leveraging BIDS specifications (***Niso et al., 2018***; ***Pernet et al., 2019***). The pipeline performed automatic bad channel detection (both noisy and flat), then applied Maxwell filtering and Signal Space Separation on the raw data (***Taulu and Kajola, 2005***). The data was then filtered between .1 Hz and 40Hz, and resampled to 250 Hz. Extraction of epochs was performed for each shape, starting 150ms before stimulus onset and stopping 1150ms after, and the relevant metadata (event type, run, trial index, etc.) for each epoch was recovered from the stimulation procedure at this step. Artifacts in the data (e.g. blinks, and heartbeats) were repaired with signal-space projection (***Uusitalo and Ilmoniemi, 1997***), and thresholds derived with “autoreject global” (***Jas et al., 2017***). For source reconstruction, some preprocessing steps were performed by fmriprep (see below). Then, sources were positioned using the “oct5” spacing with 1026 sources per hemisphere, and we used the e(xact)LORETA method (following recommendations from the literature (***Jatoi et al., 2014***; ***Pascual-Marqui et al., 2018***)) using empty-room recordings performed right before or right after the experiment to estimate the noise covariance matrix. Additionally, for source reconstruction anatomical MRI were preprocessed with fmriprep. T1-weighted (T1w) images were corrected for intensity non-uniformity (INU) with N4BiasFieldCorrection (***Tustison et al., 2010***), distributed with ANTs 2.3.3 (***Avants et al., 2008***).

The T1w-reference was then skull-stripped with a Nipype implementation of the antsBrainExtraction.sh workflow. Brain tissue segmentation of cerebrospinal fluid, white-matter and gray-matter was performed on the brain-extracted T1w using fast (***Zhang et al., 2001***). Brain surfaces were reconstructed using recon-all (***Dale et al., 1999***) and the brain mask estimated previously was refined with a custom variation of the method to reconcile ANTs-derived and FreeSurfer-derived segmentations of the cortical gray-matter of Mindboggle (***Klein et al., 2017***).

#### Decoding

After epoching the data, for each timepoint within an epoch and each participant, we trained a logistic regression decoder to classify epochs as reference or oddball using samples from all shapes. For this analysis we discarded the 6 first trials at the beginning of each block, since (i) those could not be oddballs ever and (ii) there was no warning of transitions between blocks and so the first trials were also “oddballs” with respect to the previous block’s shape. Each epoch was normalized before training the classifier. To avoid decoders being biased by the overall signal’s autocorrelation across timescales, we used six folds of cross validation over runs with the following folds: even runs versus odd runs; first half versus second half; and runs 1, 2, 5, 6 versus 3, 4, 7 and 8. Folds were used in both directions, e.g. even for training and odd for testing; as well as odd for training and even for testing. The decoders were trained on data from all of the shapes conjointly. When testing its accuracy, we tested it separately on data from each shape (e.g. detecting oddballs within squares only, within rectangles only, etc.) – using runs independent from the training data. In order to estimate accuracy without being biased by the imbalanced number of epochs in the different classes (there are 4 oddballs for every 20 references), we report the ROC Area Under the Curve of the Receiver Operating Characteristic (ROC AUC) for each shape in Fig.4B, left subfigure. Then at each timepoint, we correlated the decoding performance with the number of geometric features present in each shape: the r correlation coefficient at each timepoint is plotted in the central subfigure, together with shading for a significant cluster identified with permutation tests across participants (implemented by mne, one tailed, 2^13^ permutations, cluster forming threshold <.05). Finally, we average the decoding performance in the identified cluster and plot each shape’s average decoding performance against online behavioral data: this is effectively visualizing the same data as the central column’s figure, and therefore no statistical test is reported. The analyses were performed both without any smoothing, and with a uniform averaging over a sliding window of 100ms; the results are identical but we chose the latter since plots using the smoothed version make the separation of the different shapes easier to see. The same holds true for the next analysis.

In Fig.4C, we display a similar sequence of plots, but now instead of training a single classifier to identify epochs as reference or oddball conjointly on all shapes, we train eleven separate such classifier, one for each shape. This produces very similar results from the previous analysis.

#### RSA analysis

For RSA analyses, data from the oddballs was discarded and we used data from the magnetometers only. The goal of this analysis was to pinpoint when and where the mental representation of shapes followed distances that matched either a neural network model or a geometric feature model. We used the same model RDMs as the one we used to analyze behavior and fMRI data, and provide below the details of how we derived the empirical RDMs for our analyses.

#### In sensor space

We estimated, at each timepoint, the dissimilarity between each pair of shape across sensors: we relied on rsatoolbox to compute the Mahalanobis distance, crossvalidated with a leave-one-out scheme over runs. This provided us with one empirical RDMs for each timepoint, with no spatial information as this was performed across all sensors. Then we compared this RDM with our two models: since our model RDMs are effectively orthogonal, we performed the comparisons with the empirical RDM separately. We used a whitened correlation metric to compare RDMs. This gave use one timeseries for each participant and each model, and we then performed permutation testing in the [0, 800]ms window to identify significant temporal clusters for each model separately. As shown in Fig.4C, this yields one significant cluster associated to the CNN encoding model, and two significant cluster associated to the geometric feature model.

#### In source space

In order to understand not only the temporal dynamic of the mental representations, but also get an estimate of the localization of the various representations, we turned to RSA analysis in source space. We performed source reconstruction of each reference shape trial. Then we averaged the data over the two identified temporal clusters from the sensor-space RSA analysis (we merged the two clusters associated to the exact geometric feature model): [60, 320]ms and [128, 400]ms. We then performed RSA analysis at each source’s location using its neighboring sources (geodesic distance of 2cm on the reconstructed cortical surface). Finally, we compared the resulting RDMs to either the CNN encoding model or the geometric feature model. These steps were performed independently for each subject. Next, we projected the whitened correlation distance between empirical RDMs and model RDM onto a common cortical space, fsaverage. Finally, we performed a permutation spatial cluster test across participants, using adjacency matrices between sources (sources are adjacent if they were immediate neighbors on the reconstructed surface’s mesh). This resulted in two significant clusters associated to the CNN encoding model during the [60, 320]ms time window, located in bilateral occipital areas. Additionally, this resulted in two significant clusters associated to the geometric feature model during the [128, 400]ms, located in very broad bilateral networks encompassing dorsal and frontal areas.

## Supporting information

Video 1: RSA in source space (Exp. 4)

## Acknowledgments

We are grateful to Ghislaine Dehaene-Lambertz, Leila Azizi, and the NeuroSpin support teams for help in data acquisition, and Lorenzo Ciccione, Christophe Pallier, Minye Zhan, Alexandre Gramfort and the MNE team for support in data processing.

## Author contributions

Conceptualization: MSM, SD

Methodology: MSM, SD

Investigation: MSM, LB, CPW, CH, FAR

Visualization: MSM, LB, FAR, SD

Funding acquisition: SD

Project administration: MSM, SD

Supervision: SD

Writing – original draft: MSM, SD

Writing – review & editing: all authors

## Data and materials availability

scripts for all of the analyses are available at https://github.com/mathias-sm/AGeometricShapeRegularityEffectHumanBrain; behavioral data and scripts to generate the models are also available at this url. Raw fMRI data is provided at https://openneuro.org/datasets/ds006010/versions/1.0.1 and raw MEG data at https://openneuro.org/datasets/ds006012/versions/1.0.1

## Additional Materials and Methods

### Ethics

All studies were conducted in accordance with the Declaration of Helsinki and French bioethics laws. On-line collection of behavioral data was approved by the Paris-Saclay University Committee for Ethical Research (CER CER-Paris-Saclay-2019-063). Behavioral and brain-imaging studies in adults and 6-year-old children (MEG and fMRI) were approved by the CEA ethics boards and by a nationally approved ethics committee (MEG: CPP_100049; fMRI in children: CPP 100027 and CPP 100053; in adults CPP 100050)

## Supplementary text

### Behavioral intruder task and generation of behavioral similarity matrix Additional Results in the Behavior task: comparison with other CNNs

In Fig.S1 we report additional results from correlating human behavior with various models (top; DenseNet and ResNet) and various layers of a single model (bottom; many layers of CORnet). In all cases the CNN encoding model is a much worse predictor of the human behavior than the geometric feature model (all p<.001). The late layers of CORnet, DenseNet and ResNet all capture some of the variance of participants behaviors to comparable; while ResNet is the highest scoring in this analysis, we can see in Fig.S1A that they are all highly correlated, and in order to stay close to previous work we used CORnet’s layer IT throughout the article.

Additionally, the fit of the successive layers of CORnet (V1, V2, V4: small effect sizes, only significant for V1; IT, flatten: much higher effect sizes, increasing between IT and flatten) indicates that the feed-forward processing of the visual input by the CNN encoding yields internal representations that are increasingly closer to humans’. This suggest that even the visual model goes beyond local properties that V1 would capture – but in all cases, the fit is much worse than the geometric feature model.

### Intruder task during fMRI

#### Intruder task

Overall, both age groups performed better than chance (all p<.001) at detecting the intruder, with an average response time of 1.03s in adults, 1.36s in children, for the easy condition, and 0.79s in adults in the hard condition. Both error rates and response times are modulated by the reference shape (all p <.001), and both are correlated with error rate data from outside the scanner (all p<.05).

### Intruder task in MEG Participants

Participants recruited for the MEG studied performed no behavioral task inside the MEG, as we relied on a passive presentation oddball paradigm. After the scanner session, participants took one run of the intruder detection task previously used online and described in (***Sablé-Meyer et al., 2021***) – we presented them with the task after the scanning session to avoid biasing participants toward actively looking for intruders in the oddball paradigm. At the group level, the data fully replicated the geometric regularity effect (***Sablé-Meyer et al., 2021***) (correlation of error rate with that of previous participants, regression over 11 shapes: r^2^=.90, p<.001). A mixed effect model correlating the error rates of the two groups, with both the slope and the intercept as separate random effects, yielded an intercept not significantly different from 0 (p=.42), and a slope significantly different from 0 (p<1e-10) and not significantly different from 1 (p=.19) suggesting that the data was similar in the two groups. We also correlated each participant’s average error rate per shape to the group data from our previously dataset (***Sablé-Meyer et al., 2021***). A one-tailed test for a positive slope indicated that 19 out of 20 participants displayed a significant geometric regularity effect. We still included the data from the participant that did not display a significant effect, as we had not decided on such a rejection criterion beforehand.

### fMRI contrasts for geometric shape in the visual localizer

In order to isolate the brain responses to geometric shapes, we focused on the simplest possible contrast, i.e. greater activation to the presentation of a single geometric shape versus all of its single-image controls (face, house, tool). This contrast is presented in Figure 2C. Note that we excluded Chinese characters from this comparison because they often include geometric features (e.g. parallel lines), but including them gave virtually identical results (compare Fig.S3A and Fig.S3B, identical list of cluster-level corrected clusters at the p<.05 level in both age groups). We also included in the design rows of three distinct geometric shapes (e.g. square, triangle, circle). Our logic here was that this condition, although somewhat artificial from the geometric viewpoint, could be very tightly matched with two other conditions, namely a string of three letters (“words” condition, e.g. BON) or a small 3-symbol mathematical operation (“numbers” condition, e.g. 3+1). However, the corresponding contrast (3 geometric shapes > words and numbers) did not give any positive activation: no positive cluster was significant at the whole brain cluster-level corrected p<.05 level in either age group; additionally, none of the ROIs identified in the single shape contrast was significant for this contrast: aIPS left, p=.975 for adults and p=.389 for children; aIPS right, p=.09 in both adults and children; pITG right, p=.13 in adults and p=.362 in children). We reasoned, however, that numbers should be excluded from this contrast because, by our very hypothesis, geometric shapes should activate a number-based program-like representations (e.g. square = repeat(4){line, turn}) (***Sablé-Meyer et al., 2022***). When restricting the contrast to “3 geometric shapes > words”, at the whole brain level, no cluster reached significance at the p<.05 level (cluster-level correction). However, in this case, testing the ROIs identified in the single shape condition revealed that while the left aIPS still did not reach significance at the p<.05 level (p=.71 in adults, p=.30 in children), both the right aIPS and the right pITG reached significance (aIPS right: p<.001 in adults and p=.014 in chidren; pITG, p=.008 in adults and p=.046 in children).

### Additional ROI analysis of the ventral pathway in fMRI (Fig.S4)

We used individually defined regions of interest (ROIs) to probe the fMRI response to geometric shapes in four cortical regions defined by their preferential responses to four other visual categories known to be selectively processed in the ventral visual pathway: faces, houses, tools and words. To this aim, we first identified, at the group level, clusters of voxels associated to each visual category. Such clusters were identified using a non-parametric permutation test across participants at the whole-brain level using a contrast for the target category against the mean of the other three (voxel p<.001 then clusters p<.05 except for ffa in children, p=.09, and the absence of a well-identified vwfa in children). Significant clusters which intersect the MNI coordinate z=-14 are shown in Fig.S4; in the case of the fusiform face area (FFA) and the visual word form area (VWFA), we restricted ourselves to clusters at MNI coordinates typically found in the literature, respectively (−45, −57, −14) and (40, −55, −14).

Then, within each such ROI identified in adults, we identified for each subject, including children, the 10% most responsive subject-specific voxels in the same contrast used to identify the cluster. To avoid double-dipping, we selected the voxels using the contrast from a single run, then collected the fMRI responses (beta coefficients) to all categories from the other runs, and then replicated this procedure across all runs while averaging the responses to a given category. The average coefficients within each such individually-defined cortical ROI are shown in Fig.S4, separately for children and adults. Several observations are in order, and detailed below for each visual category. In the left-hemispheric VWFA, we can see that voxels are indeed responsive to written words in the participants’ language (French), more than to an unknown language (Chinese), in both adults and children (paired t-tests, p<.001 in adults, p=.003 in children). VWFA voxels also responded to the symbolic display of numbers entering into small computations (e.g. 3+1) in adults, but this response did not appear to be developed yet in children. VWFA voxels also showed a response to tools, particularly in young children, as already been reported (***Dehaene-Lambertz et al., 2018***). In the opposite direction, they were particularly under-activated by houses in adults. Finally, and most crucially, geometric shapes, whether presented alone or in a string of 3, did not elicit a strong response, indeed no stronger than non-preferred categories such as Chinese characters or faces (p=.37 in adults, p=.057 in children on a one-tailed paired t-test).

In the right-hemispheric FFA, we only saw a purely selective response to faces in both adults and children. All the other visual categories yielded an equally low level of activity in this area. In particular, the responses to geometric shapes were not significantly different from those to other visual categories.

In the bilateral ROIs responsive to tools, a very similar result was found: apart from their strong response to tools and their slightly increased activation to houses and reduced activations to written words and numbers in the right hemisphere, these voxels elicit activations that were essentially equal across all non-preferred visual categories, with geometric shapes being no exception.

Finally, bilateral house-responsive ROIs corresponding to the parahippocampal place area (PPA) were also mildly responsive to tools in both populations and both hemispheres. However, geometric shapes did not activate these clusters more than other visual categories such as faces (both hemispheres, both age groups have p>.05 in one-tailed paired t-tests).

### Additional analysis: Evoked Reponse Potentials (ERP)

Different participants can in principle end up with very different decoder weights, since the decoders are trained independently for each participant. Several of our analyses are based on the decoders’ performance only, and therefore the decoder’s weights associated to each sensor are not considered. To stay closer to the MEG data, we replicated the previous analysis directly from evoked potential data in the gradiometers. Using spatiotemporal permutation testing, we identified a set of sensors ant timepoints across participants where the reference and the oddball epochs were significantly different. We identified three significant clusters: [268, 844]ms, [440, 896]ms and [492, 896]ms.

Then we computed the average difference between the reference and the oddball trials across these sensors separately for each shape. Finally, we correlated the differences with the number of geometric features in the reference shape. The first cluster, from 268 ms to 844 ms, did not elicit any significant correlation with the geometric regularity; however, the two others yielded significant clusters at the p<.05 level, although with later timing than the decoding analysis, at around ∼600ms.

### MEG: Joint MEG-fMRI RSA

Representational similarity analysis also offers a way to directly compared similarity matrices measured in MEG and fMRI, thus allowing for fusion of those two modalities and tentatively assigning a “time stamp” to distinct MRI clusters (***Cichy and Oliva, 2020***). However, we did not attempt such an analysis here for several reasons. First, distinct tasks and block structures were used in MEG and fMRI. Second, a smaller list of shapes was used in fMRI, as imposed by the slower modality of acquisition. Third, our study was designed as an attempt to sort out between two models of geometric shape recognition. We therefore focused all analyses on this goal, which could not have been achieved by direct MEG-fMRI fusion, but required correlation with independently obtained model predictions.

**Fig. S1.**
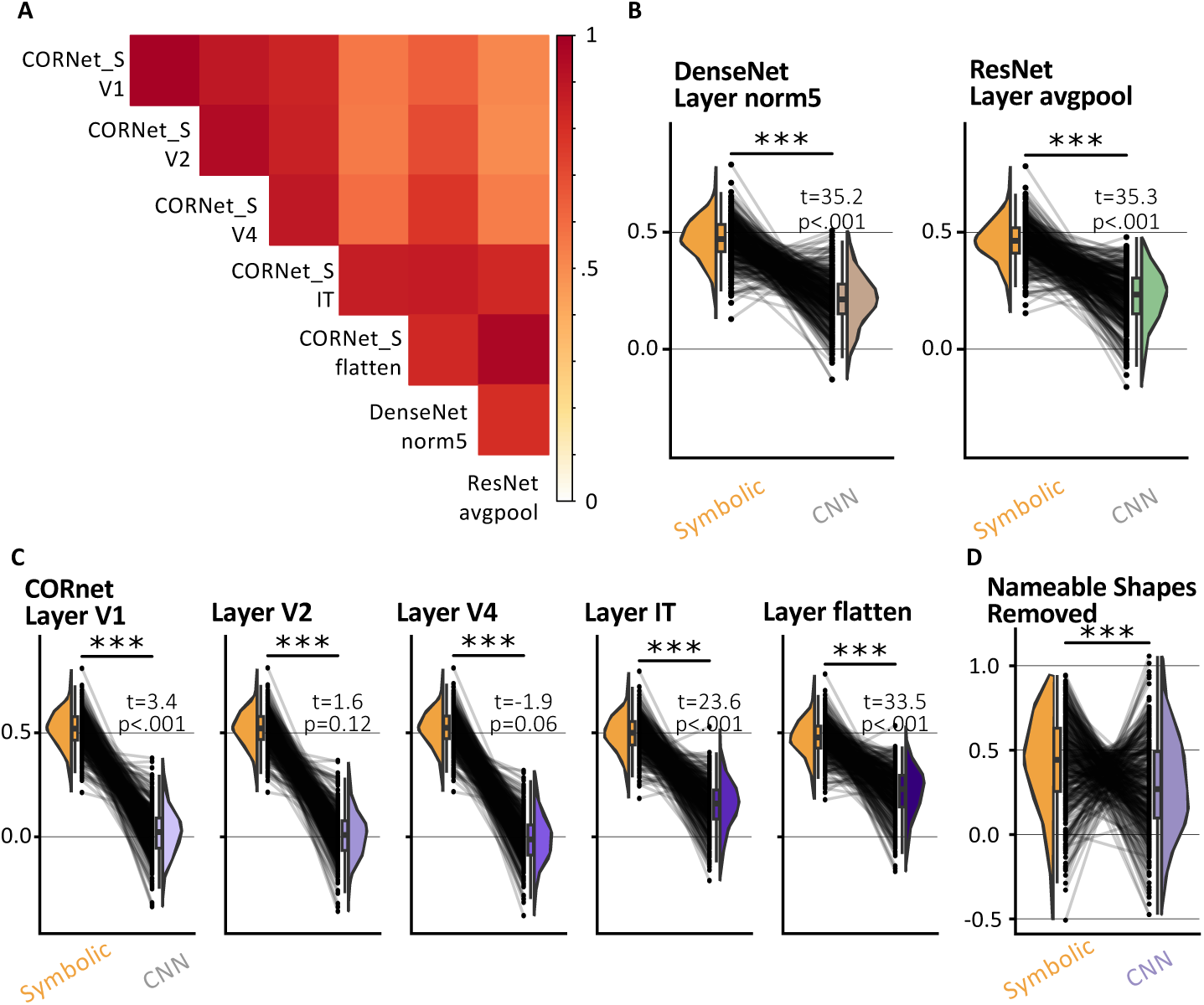
Additional CNN encoding models of human behavior. **A**, Correlation matrix between all pairs of RDMs generated by each CNN and layer considered. **B**, Replication of Fig.1D with different CNNs. Stars indicate a significant difference between the geometric feature model and the respective CNN encoding model (p<0.001). t and p values also indicate whether the CNN encoding model is a significant predictor of participant’s behavior (the geometric feature model is always highly significant). **C** Replication of B using different layers of CORnet, organized from early to late layers, from left to right. Note that the late layers are much more significant predictors of human behavior than the early ones – although still far inferior to the geometric feature model. **D** Replication of our GLM analysis including only shapes for which there is no obvious name in English–though we gave them names in this manuscriptto refer to them: “kite”, “rightKite”, “hinge”, “rustedHinge” and “random”.

## Figures

### Video

Video 1: searchlight based timepoint-per-timepoint RSA analysis across shapes. Significant (p<.05) sources associated to the CNN model are shown in purple, significant sources associated to the Geometric Feature model in orange, and associated to both in green.

### Tables

**Table S1. Coordinates and characteristics of significant fMRI clusters responding to geometric shapes in localizer runs**

**Table S2. Coordinates and characteristics of significant fMRI clusters in the RSA analysis**

**Fig. S2.**
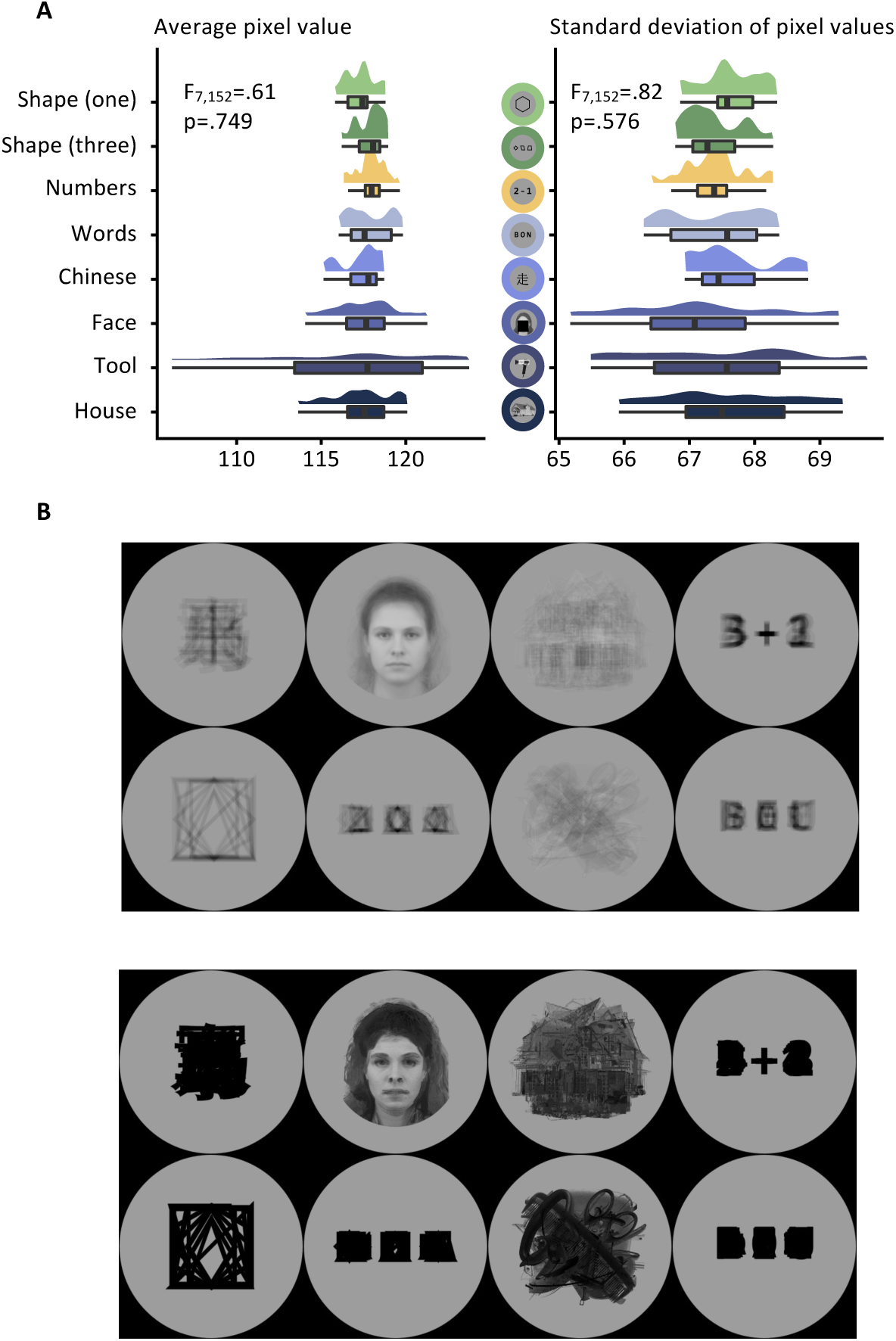
Overview of the stimuli used for the category localizer. **A**, Average pixel value (left) and average standard deviation across pixels (right) for stimuli within each category (y axis). An ANOVA indicated no significant effect of the stimulus category on either the average or the standard deviation across pixels. **B**, Average (top) and max (bottom) pixel value at each location across the eight possible visual categories used in the localizer.

**Fig. S3.**
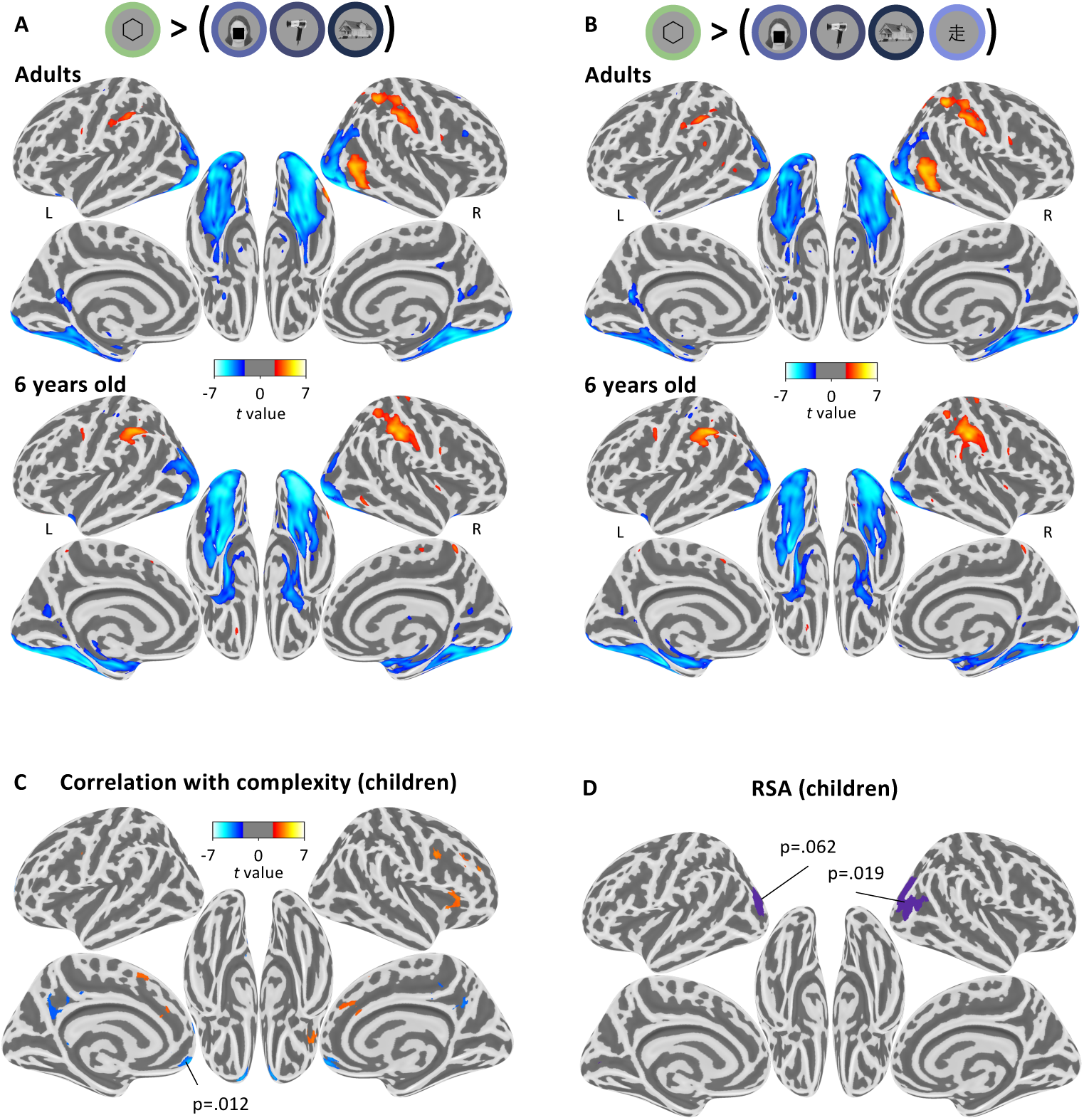
Details of the fMRI results in children (complement to Fig.2 and Fig.3 in the main text) **A:** Statistical map associated with the contrast “single geometric shape > faces, houses and tools”, projected on an inflated brain (top: adults; bottom: children; for illustration purpose we display the uncorrected statistical map at the p<.01 level). Notice how similar the activations are in both age groups. **B:** Same as A, but for the contrast “single geometric shape > all single-object visual categories (face, house, tools, Chinese characters)”. The activation maps are very similar to the previous contrast, and very similar across age groups. **C:** Whole brain correlation of the BOLD signal with geometric regularity in children, as measured by the error rate in a previous online intruder detection task (***Sablé-Meyer et al., 2021***). Positive correlations are shown in red and negative ones in blue. Voxel threshold p<.001, no correction for multiple comparisons, but the p-value indicates the only cluster that was significant at the cluster-level corrected p<.05 threshold. **D,** Results of RSA analysis in children. No cluster was significant at the p<.05 level for the geometric feature models; one right-lateralized occipital cluster reached significance for the CNN encoding model (cluster-level corrected p=.019), and its symmetrical counterpart was close to the significance threshold (cluster-level corrected p=.062)

**Fig. S4.**
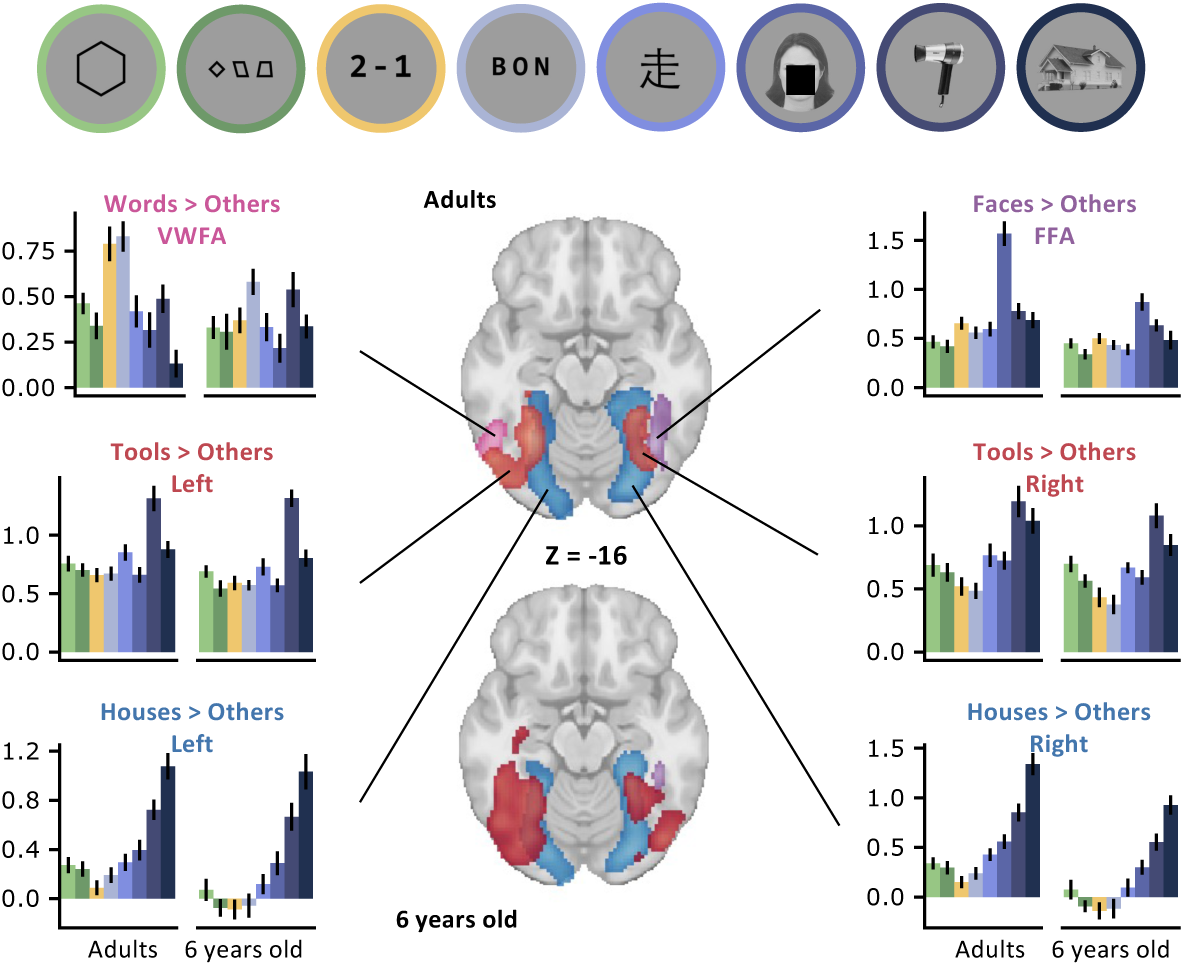
fMRI response of subject-specific voxels in the ventral visual pathway to geometric shapes and other visual stimuli. The brain slices show the group-level clusters associated to various contrasts known to elicit a selective response in the ventral visual pathway, in both age groups: VWFA (words > others; green), FFA (faces > others; purple), tool-selective ROIs (tools > others; red) and PPA (houses > others; light blue). Within each ROI, plots show the mean beta coefficients for the BOLD effect within a subject-specific selection of the 10% best voxels, using independent runs for selection and plotting to avoid circularity and “double-dipping”.

**Appendix 0—table 1.** Coordinates and characteristics of significant fMRI clusters responding to geometric shapes in localizer runs. For each age group, each line gives the peak coordinates, volume and statistics of a cluster with p<.05 (whole brain, permutation test) for the contrast “single shape > other single visual categories”. The sign of the peak t-value and the shading indicate whether the contrast was positive (white background) or negative (grey background). Coordinates are given in MNI space.

| Age Group | X | Y | Z | Peak t-value | Volume in cm<sup>2</sup> | Cluster corrected p-value |
| --- | --- | --- | --- | --- | --- | --- |
| Adults | 51.5 | -54.5 | -8.5 | 5.12 | 4.4 | p<.01 |
|  | 35.5 | -48.5 | 55.5 | 4.93 | 7.2 | p<.01 |
|  | 39.5 | -76.5 | -12.5 | -6.33 | 51.4 | p<.01 |
|  | -34.5 | -70.5 | -10.5 | -5.92 | 42 | p<.01 |
| Children | -12.5 | -76.5 | -48.5 | 5.51 | 3.9 | p=.01 |
|  | 55.5 | -24.5 | 45.5 | 5 | 8.3 | p<.01 |
|  | 21.5 | -80.5 | -46.5 | 4.9 | 4.4 | p=.01 |
|  | -42.5 | -38.5 | 35.5 | 4.9 | 6.4 | p<.01 |
|  | 21.5 | -94.5 | -8.5 | -6.34 | 40.1 | p<.01 |
|  | -24.5 | -98.5 | -8.5 | -6.17 | 55.3 | p<.01 |

**Fig. S5.**
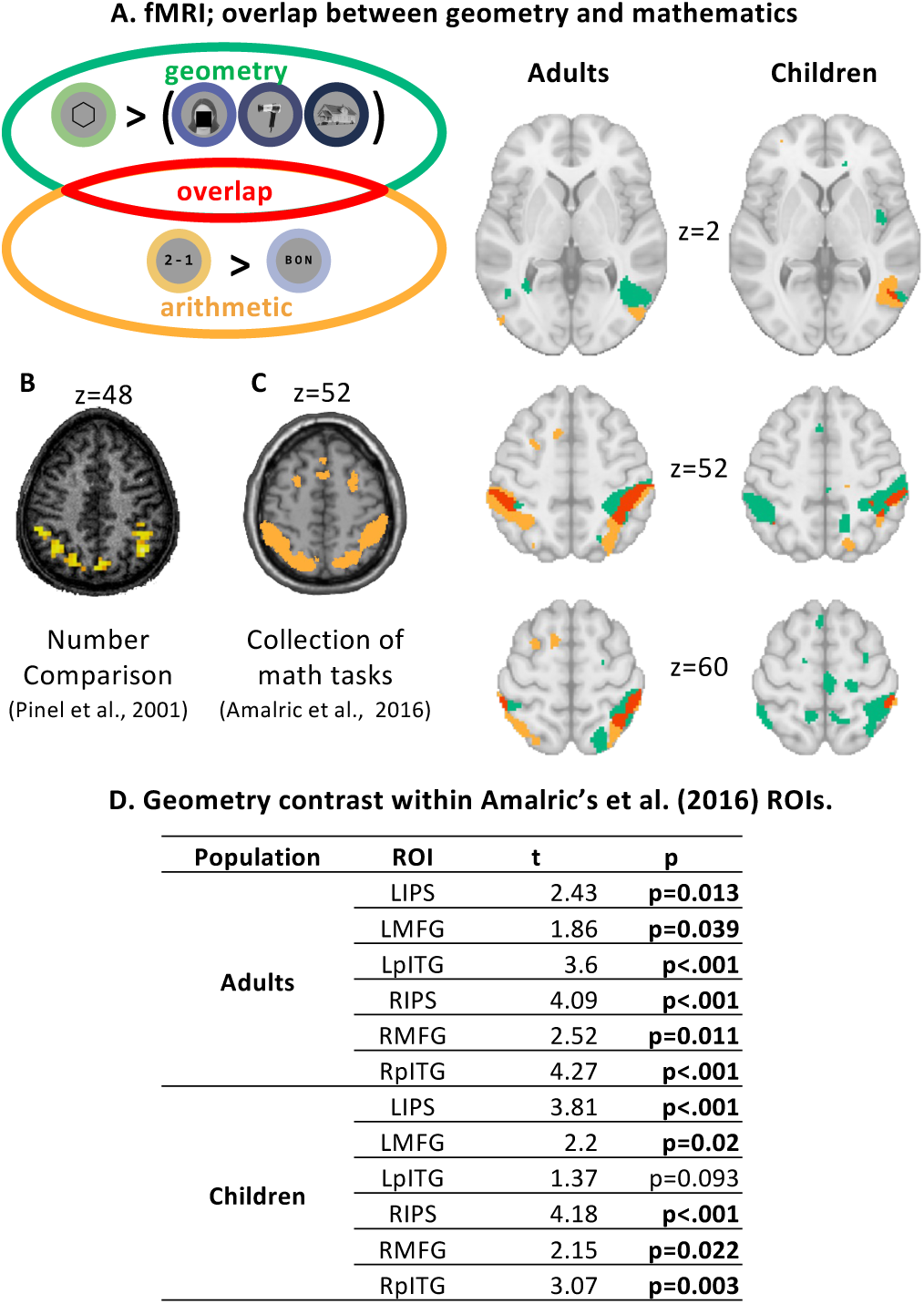
**A** Overlap in red between out geometry contrast in green (shape versus other single objects) and our number contrast in orange (numbers > words), in three slices: two that intersect with the bilateral IPS areas (z=60 and z=52) and the rITG (z=2) in both populations. To help visualize areas that coincide between populations but did not reach significance in one or the other, the maps here are uncorrected, p<.01. **B** Statistical map from (***Pinel et al., 2001***) showing areas where activation was correlated with numerical distance in a number comparison task, slice at z=48 (p<.001, uncorrected). **C** Statistical map from (***Amalric and Dehaene, 2016***) showing the overlap between three math-related tasks, including high-level mathematical judgments in mathematicians; slice at z=52. **D** Statistical tests for the “shape > other categories” contrast from the ROIs identified in independent work (***Amalric and Dehaene, 2016***) in both populations, with all ROIs having a significant test at the p<.05 level except the LpITG.

**Appendix 0—table 2.**
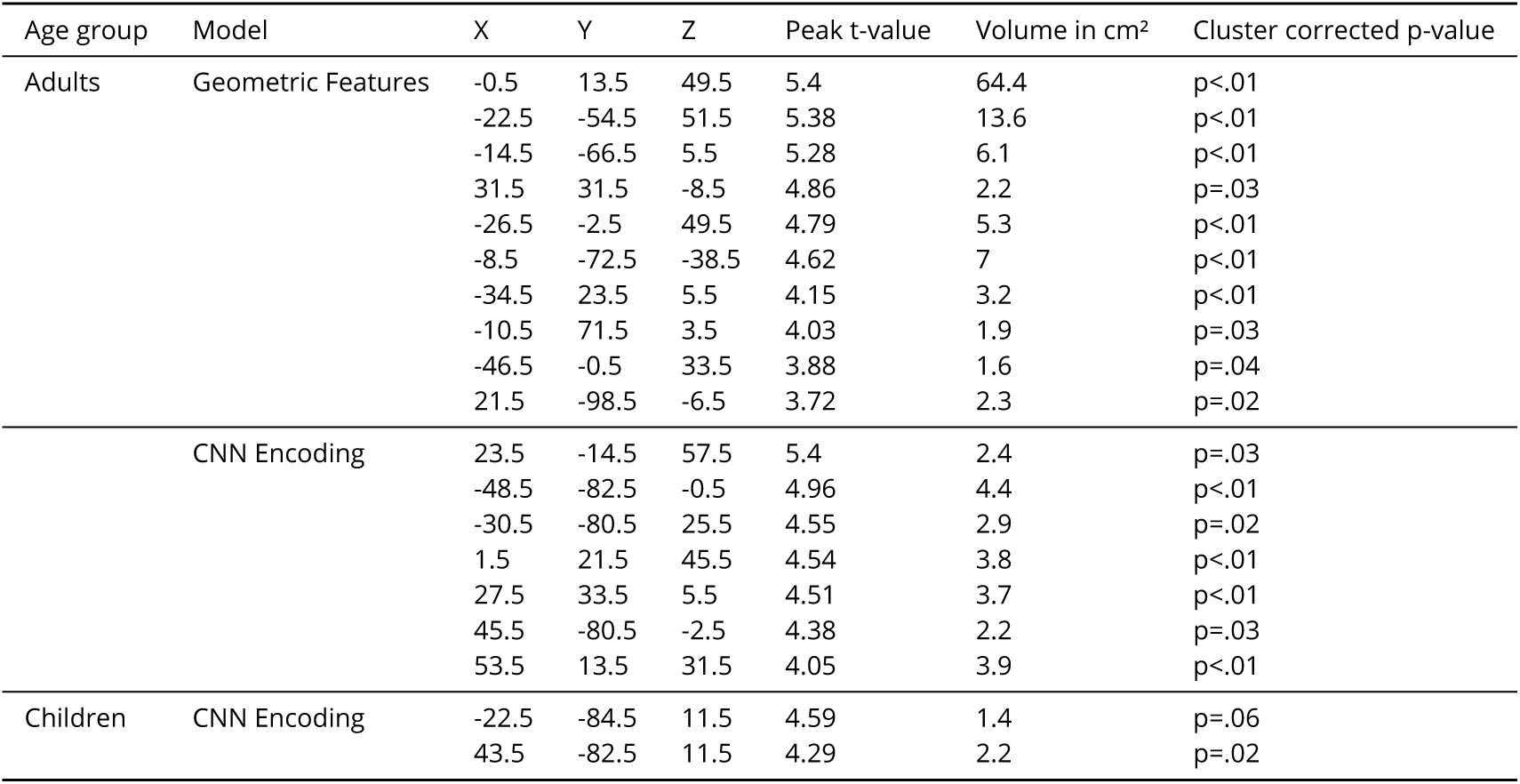
Coordinates and characteristics of significant fMRI clusters in the RSA analysis. Same organisation as table S1 for the RSA analysis.

**Fig. S6.**
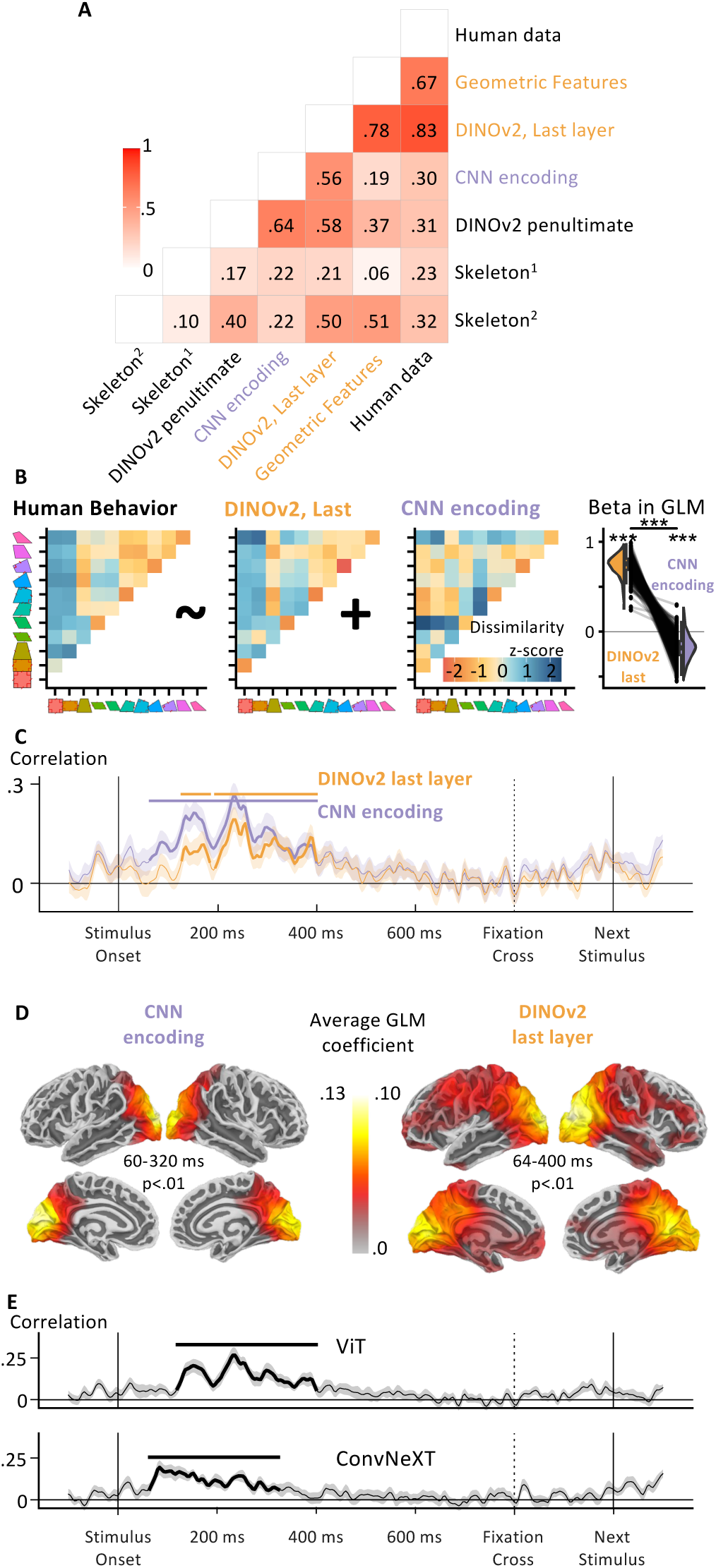
Additional Models: Behavior and MEG. **A.** Correlation between several models and the average RDM across participants. In particular, we have added the last two layers of DINOv2 (***Oquab et al., 2023***) as well as two different implementation of distances in skeletal spaces (***Ayzenberg and Lourenco, 2019***; ***Morfoisse and Izard, 2021***) **B.** Fig. 1D with the symbolic model replaced with the empirical RDM obtained from the last layer of DINOv2 using Euclidean distance. **C.** Fig. 4C with the same replacement of the symbolic model with the last layer of DINOv2. **D.** Fig. 4D with the same replacement of the symbolic model with the last layer of DINOv2. E. Timecourse of the similarity between empirical RDMs and two additional neural networks: a vision transformer network (ViT, top (***Dosovitskiy et al., 2020***)), and a large CNN (ConvNeXT, bottom, (***Liu et al., 2022***)), both with many parameters (respectively ∼1 billion and ∼800 million) and trained on 2 billion images (***Cherti et al., 2023***; ***Schuhmann et al., 2022***)).

**Fig. S7.**
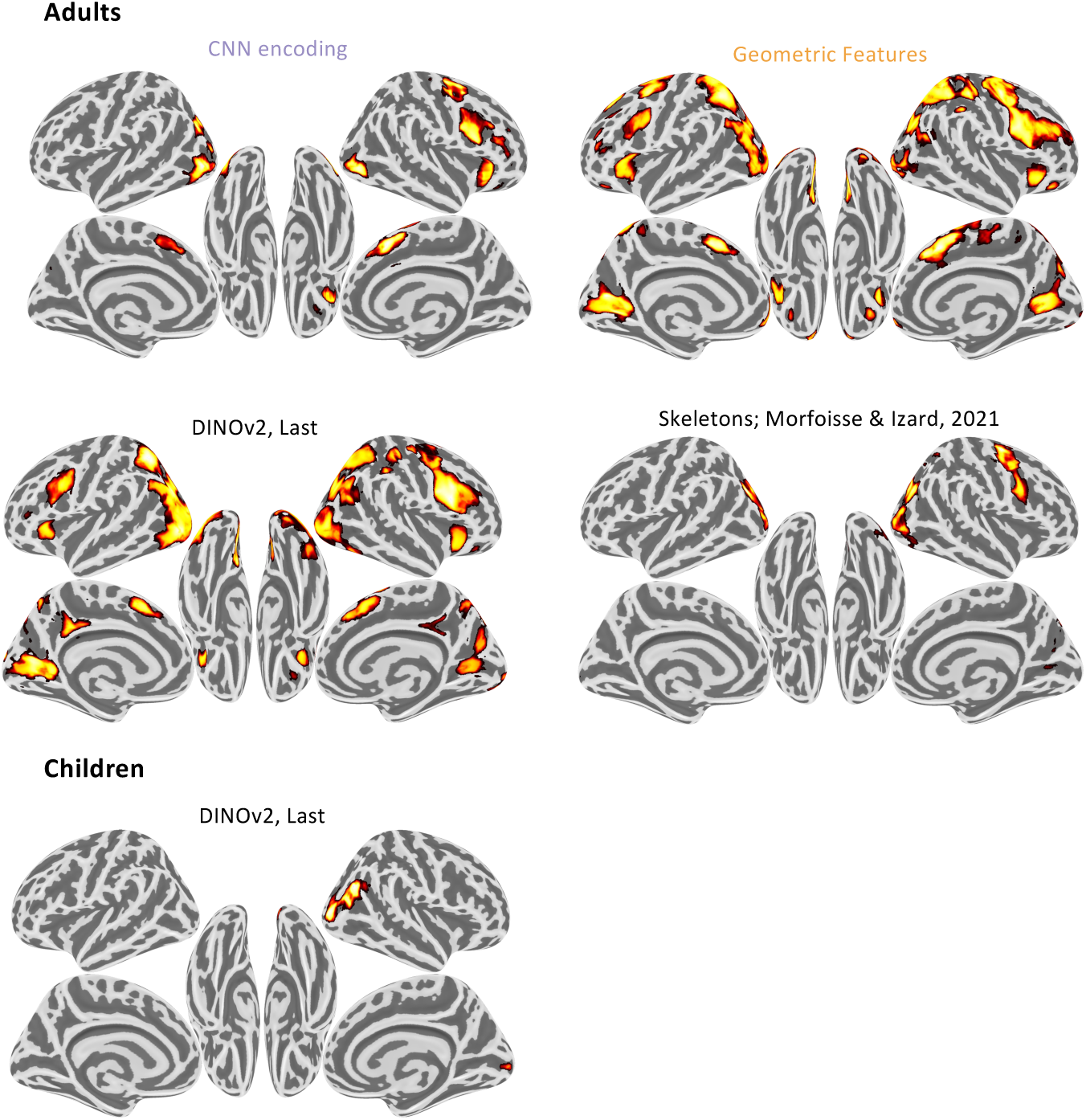
Additional Models: fMRI. t-values inside significant clusters at the p<.05 level for four models: geometric features (top left), CNN encoding (top right), DINOv2 last layer (bottom left) and skeletal representations from (***Morfoisse and Izard, 2021***) (bottom right). Skeletal representations from (***Ayzenberg and Lourenco, 2019***) did not yield any significant clusters in adults. In children (bottom), only the DINOv2 elicited any significant cluster.

## References

Abraham A, Pedregosa F, Eickenberg M, Gervais P, Mueller A, Kossaifi J, Gramfort A, Thirion B, Varoquaux G. Machine learning for neuroimaging with scikit-learn. Frontiers in Neuroinformatics. 2014; 8. https://www.frontiersin.org/articles/10.3389/fninf.2014.00014/full, doi: 10.3389/fninf.2014.00014.

Agrawal A, Hari K, Arun S. Reading Increases the Compositionality of Visual Word Representations. Psychological Science. 2019; 30(12):1707–1723.

Agrawal A, Hari K, Arun S. A Compositional Neural Code in High-Level Visual Cortex Can Explain Jumbled Word Reading. eLife. 2020; 9:e54846. doi: 10.7554/eLife.54846.

Al Roumi F, Planton S, Wang L, Dehaene S. Compression of Binary Sound Sequences in Human Memory. eLife. 2023; in press:2022.10.15.512361. https://www.biorxiv.org/content/10.1101/2022.10.15.512361v1.

Allen EJ, St-Yves G, Wu Y, Breedlove JL, Prince JS, Dowdle LT, Nau M, Caron B, Pestilli F, Charest I, Hutchinson JB, Naselaris T, Kay K. A Massive 7T fMRI Dataset to Bridge Cognitive Neuroscience and Artificial Intelligence. Nature Neuroscience. 2021; p. 1–11. doi: 10.1038/s41593-021-00962-x, bandiera_abtest: a Cg_type: Nature Research Journals Primary_atype: Research publisher: Nature Publishing Group Subject_term: Cortex;Neural encoding;Object vision;Perception Subject_term_id: cortex;neural-encoding;object-vision;perception.

Amalric M, Dehaene S. Origins of the Brain Networks for Advanced Mathematics in Expert Mathematicians. Proceedings of the National Academy of Sciences. 2016; p. 201603205. doi: 10.1073/pnas.1603205113.

Amalric M, Dehaene S. A Distinct Cortical Network for Mathematical Knowledge in the Human Brain. NeuroImage. 2019; 189:19–31.

Amalric M, Denghien I, Dehaene S. On the Role of Visual Experience in Mathematical Development: Evidence from Blind Mathematicians. Developmental Cognitive Neuroscience. 2018; 30:314–323. doi: 10.1016/j.dcn.2017.09.007.

Amir O, Biederman I, Hayworth KJ. Sensitivity to Nonaccidental Properties Across Various Shape Dimensions. Vision Research. 2012; 62:35–43. doi: 10.1016/j.visres.2012.03.020.

Ansari D. Effects of Development and Enculturation on Number Representation in the Brain. Nat Rev Neurosci. 2008; 9(4):278–91.

Arcaro MJ, Schade PF, Vincent JL, Ponce CR, Livingstone MS. Seeing Faces Is Necessary for Face-Domain Formation. Nature Neuroscience. 2017; 20(10):1404–1412. doi: 10.1038/nn.4635.

Avants BB, Epstein CL, Grossman M, Gee JC. Symmetric Diffeomorphic Image Registration with Cross-Correlation: Evaluating Automated Labeling of Elderly and Neurodegenerative Brain. Medical Image Analysis. 2008; 12(1):26–41. doi: 10.1016/j.media.2007.06.004.

Ayzenberg V, Behrmann M. Does the Brain’s Ventral Visual Pathway Compute Object Shape? Trends in Cognitive Sciences. 2022; 26(12):1119–1132. doi: 10.1016/j.tics.2022.09.019.

Ayzenberg V, Kamps FS, Dilks DD, Lourenco SF. Skeletal Representations of Shape in the Human Visual Cortex. Neuropsychologia. 2022; 164:108092.

Ayzenberg V, Lourenco SF. Skeletal Descriptions of Shape Provide Unique Perceptual Information for Object Recognition. Scientific Reports. 2019; 9(1):9359. doi: 10.1038/s41598-019-45268-y.

Ayzenberg V, Simmons C, Behrmann M. Temporal Asymmetries and Interactions Between Dorsal and Ventral Visual Pathways During Object Recognition. Cerebral Cortex Communications. 2023; 4(1):tgad003.

Bao P, She L, McGill M, Tsao DY. A Map of Object Space in Primate Inferotemporal Cortex. Nature. 2020; p. 1–6. doi: 10.1038/s41586-020-2350-5.

Behzadi Y, Restom K, Liau J, Liu TT. A Component Based Noise Correction Method (CompCor) for BOLD and Perfusion Based fMRI. NeuroImage. 2007; 37(1):90–101. doi: 10.1016/j.neuroimage.2007.04.042.

Benjamin L, Sable-Meyer M, Flo A, Dehaene-Lambertz G, Al Roumi F. Long-Horizon Associative Learning Explains Human Sensitivity to Statistical and Network Structures in Auditory Sequences. Journal of Neuroscience. 2024;.

Biederman I. Recognition-by-Components: A Theory of Human Image Understanding. Psychological review. 1987; 94(2):115.

Biederman I, Ju G. Surface Versus Edge-Based Determinants of Visual Recognition. Cognitive Psychology. 1988; 20(1):38–64. doi: 10.1016/0010-0285(88)90024-2.

Biederman I, Yue X, Davidoff J. Representation of Shape in Individuals From a Culture With Minimal Exposure to Regular, Simple Artifacts: Sensitivity to Nonaccidental Versus Metric Properties. Psychological Science. 2009; 20(12):1437–1442. doi: 10.1111/j.1467-9280.2009.02465.x.

Blum H. Biological Shape and Visual Science (Part I). Journal of Theoretical Biology. 1973; 38(2):205–287. doi: 10.1016/0022-5193(73)90175-6.

Bowers JS, Malhotra G, Dujmović M, Montero ML, Tsvetkov C, Biscione V, Puebla G, Adolfi F, Hummel JE, Heaton RF, Evans BD, Mitchell J, Blything R. Deep Problems with Neural Network Models of Human Vision. Behavioral and Brain Sciences. 2022; p. 1–74. doi: 10.1017/S0140525X22002813.

Cai Q, Brysbaert M. SUBTLEX-CH: Chinese Word and Character Frequencies Based on Film Subtitles. PloS one. 2010; 5(6):e10729.

Campbell DI, Kumar S, Giallanza T, Cohen J, Griffiths T. Human-Like Geometric Abstraction in Large Pre-trained Neural Networks. In: Proceedings of the Annual Meeting of the Cognitive Science Society, vol. 46; 2024. .

Cantlon JF, Li R. Neural activity during natural viewing of Sesame Street statistically predicts test scores in early childhood. PLoS biology. 2013; 11(1):e1001462. doi: 10.1371/journal.pbio.1001462.

Cavanagh P. The Language of Vision. Perception. 2021; 50(3):195–215. doi: 10.1177/0301006621991491.

Chater N, Vitányi P. Simplicity: a unifying principle in cognitive science? Trends in Cognitive Sciences. 2003; 7(1):19–22.

Cherti M, Beaumont R, Wightman R, Wortsman M, Ilharco G, Gordon C, Schuhmann C, Schmidt L, Jitsev J. Reproducible Scaling Laws for Contrastive Language-Image Learning. In: Proceedings of the IEEE/CVF conference on computer vision and pattern recognition; 2023. p. 2818–2829.

Cichy RM, Oliva A. A M/EEG-fMRI Fusion Primer: Resolving Human Brain Responses in Space and Time. Neuron. 2020; 0(0). https://www.cell.com/neuron/abstract/S0896-6273(20)30518-3, doi: 10.1016/j.neuron.2020.07.001.

Close J, Call J. From Colour Photographs to Black-and-White Line Drawings: An Assessment of chimpanzees’(Pan Troglodytes’) Transfer Behaviour. Animal Cognition. 2015; 18:437–449.

Collins E, Freud E, Kainerstorfer JM, Cao J, Behrmann M. Temporal Dynamics of Shape Processing Differentiate Contributions of Dorsal and Ventral Visual Pathways. Journal of Cognitive Neuroscience. 2019; 31(6):821–836.

Conwell C, Prince JS, Alvarez GA, Konkle T. What Can 5.17 Billion Regression Fits Tell Us about Artificial Models of the Human Visual System? In: SVRHM 2021 Workshop@ NeurIPS; 2021. .

Cox RW, Hyde JS. Software Tools for Analysis and Visualization of fMRI Data. NMR in Biomedicine. 1997; 10(4-5):171–178. doi: 10.1002/(SICI)1099-1492(199706/08)10:4/5<171::AID-NBM453>3.0.CO;2-L.

Dale AM, Fischl B, Sereno MI. Cortical Surface-Based Analysis: I. Segmentation and Surface Reconstruction. NeuroImage. 1999; 9(2):179–194. doi: 10.1006/nimg.1998.0395.

De Leeuw J, Mair P. Multidimensional Scaling Using Majorization: SMACOF in R. Journal of statistical software. 2009; 31:1–30.

Dehaene S, Izard V, Pica P, Spelke E. Core Knowledge of Geometry in an Amazonian Indigene Group. Science. 2006; 311:381–384.

Dehaene S. The Lascaux Rectangle : How Homo Sapiens Invented Geometry. MIT Press; 2026.

Dehaene S, Al Roumi F, Lakretz Y, Planton S, Sablé-Meyer M. Symbols and Mental Programs: A Hypothesis about Human Singularity. Trends in Cognitive Sciences. 2022; 26(9):751–766. doi: 10.1016/j.tics.2022.06.010.

Dehaene S, Izard V, Pica P, Spelke E. Core Knowledge of Geometry in an Amazonian Indigene Group. . 2006; 311:5.

Dehaene-Lambertz G, Monzalvo K, Dehaene S. The Emergence of the Visual Word Form: Longitudinal Evolution of Category-Specific Ventral Visual Areas During Reading Acquisition. PLOS Biology. 2018; 16(3):e2004103. doi: 10.1371/journal.pbio.2004103.

Denisova K, Feldman J, Su X, Singh M. Investigating Shape Representation Using Sensitivity to Part-and Axis-Based Transformations. Vision research. 2016; 126:347–361.

Diedrichsen J, Berlot E, Mur M, Schütt HH, Shahbazi M, Kriegeskorte N, Comparing Representational Geometries Using Whitened Unbiased-Distance-Matrix Similarity. arXiv; 2021. http://arxiv.org/abs/2007.02789.

Dillon MR, Duyck M, Dehaene S, Amalric M, Izard V. Geometric Categories in Cognition. Journal of Experimental Psychology: Human Perception and Performance. 2019; 45(9):1236–1247. doi: 10.1037/xhp0000663.

Dosovitskiy A, Beyer L, Kolesnikov A, Weissenborn D, Zhai X, Unterthiner T, Dehghani M, Minderer M, Heigold G, Gelly S, others. An Image Is Worth 16×16 Words: Transformers for Image Recognition at Scale. arXiv preprint arXiv:201011929. 2020;.

Emberson LL, Crosswhite SL, Richards JE, Aslin RN. The Lateral Occipital Cortex Is Selective for Object Shape, Not Texture/Color, at Six Months. Journal of neuroscience. 2017; 37(13):3698–3703.

Epstein R, Harris A, Stanley D, Kanwisher N. The Parahippocampal Place Area: Recognition, Navigation, or Encoding? Neuron. 1999; doi: 10.1016/S0896-6273(00)80758-8.

Esteban O, Blair R, Markiewicz CJ, Berleant SL, Moodie C, Ma F, Isik AI, Erramuzpe A, Kent James D and Goncalves M, DuPre E, Sitek KR, Gomez DEP, Lurie DJ, Ye Z, Poldrack RA, Gorgolewski KJ. fMRIPrep. Software. 2018; doi: 10.5281/zenodo.852659.

Esteban O, Markiewicz C, Blair RW, Moodie C, Isik AI, Erramuzpe Aliaga A, Kent J, Goncalves M, DuPre E, Snyder M, Oya H, Ghosh S, Wright J, Durnez J, Poldrack R, Gorgolewski KJ. fMRIPrep: A Robust Preprocessing Pipeline for Functional MRI. Nature Methods. 2018; doi: 10.1038/s41592-018-0235-4.

Evans A, Janke A, Collins D, Baillet S. Brain Templates and Atlases. NeuroImage. 2012; 62(2):911–922. doi: 10.1016/j.neuroimage.2012.01.024.

Feldman J. The Simplicity Principle in Human Concept Learning. Current Directions in Psychological Science (Wiley-Blackwell). 2003; 12(6):227–232. doi: 10.1046/j.0963-7214.2003.01267.x.

Feldman J, Singh M. Bayesian Estimation of the Shape Skeleton. Proceedings of the National Academy of Sciences. 2006; 103(47):18014–18019.

Firestone C, Scholl BJ. “Please Tap the Shape, Anywhere You Like” Shape Skeletons in Human Vision Revealed by an Exceedingly Simple Measure. Psychological science. 2014; 25(2):377–386.

Fodor JA. The Language of Thought, vol. 5. Harvard University Press; 1975.

Fonov V, Evans A, McKinstry R, Almli C, Collins D. Unbiased Nonlinear Average Age-Appropriate Brain Templates from Birth to Adulthood. NeuroImage. 2009; 47, Supplement 1:S102. doi: 10.1016/S1053-8119(09)70884-5.

Freud E, Plaut DC, Behrmann M. ‘What’is Happening in the Dorsal Visual Pathway. Trends in cognitive sciences. 2016; 20(10):773–784.

Froyen V, Feldman J, Singh M. Bayesian Hierarchical Grouping: Perceptual Grouping as Mixture Estimation. Psychological Review. 2015; 122(4):575–597. doi: 10.1037/a0039540.

Goodenough FL. Studies in the Psychology of Children’s Drawings. Psychological Bulletin. 1928; 25(5):272–283. doi: 10.1037/h0071049.

Gorgolewski K, Burns CD, Madison C, Clark D, Halchenko YO, Waskom ML, Ghosh S. Nipype: A Flexible, Lightweight and Extensible Neuroimaging Data Processing Framework in Python. Frontiers in Neuroinformatics. 2011; 5:13. doi: 10.3389/fninf.2011.00013.

Gorgolewski KJ, Esteban O, Markiewicz CJ, Ziegler E, Ellis DG, Notter MP, Jarecka D, Johnson H, Burns C, Manhães-Savio A, Hamalainen C, Yvernault B, Salo T, Jordan K, Goncalves M, Waskom M, Clark D, Wong J, Loney F, Modat M, et al. Nipype. Software. 2018; doi: 10.5281/zenodo.596855.

Greve DN, Fischl B. Accurate and Robust Brain Image Alignment Using Boundary-Based Registration. NeuroImage. 2009; 48(1):63–72. doi: 10.1016/j.neuroimage.2009.06.060.

Grill-Spector K, Kourtzi Z, Kanwisher N. The Lateral Occipital Complex and Its Role in Object Recognition. Vision research. 2001; 41(10-11):1409–1422.

Grill-Spector K, Kushnir T, Hendler T, Edelman S, Itzchak Y, Malach R. A Sequence of Object-Processing Stages Revealed by fMRI in the Human Occipital Lobe. Human brain mapping. 1998; 6(4):316–328.

He K, Zhang X, Ren S, Sun J. Deep Residual Learning for Image Recognition. In: Proceedings of the IEEE conference on computer vision and pattern recognition; 2016. p. 770–778.

Heinke D, Wachman P, van Zoest W, Leek EC. A Failure to Learn Object Shape Geometry: Implications for Convolutional Neural Networks as Plausible Models of Biological Vision. Vision Research. 2021; 189:81–92. doi: 10.1016/j.visres.2021.09.004.

Henshilwood CS, d’ Errico F, van Niekerk KL, Dayet L, Queffelec A, Pollarolo L. An Abstract Drawing from the 73,000-Year-Old Levels at Blombos Cave, South Africa. Nature. 2018; 562(7725):115–118. doi: 10.1038/s41586-018-0514-3.

Huang G, Liu Z, van der Maaten L, Weinberger KQ. Densely Connected Convolutional Networks. arXiv:160806993 [cs]. 2018; http://arxiv.org/abs/1608.06993.

Izard V, Dehaene-Lambertz G, Dehaene S. Distinct Cerebral Pathways for Object Identity and Number in Human Infants. PLoS Biol. 2008; 6(2):275–285.

Izard V, Pica P, Spelke ES. Visual Foundations of Euclidean Geometry. Cognitive Psychology. 2022; 136:101494. doi: 10.1016/j.cogpsych.2022.101494.

Izard V, Pica P, Spelke ES, Dehaene S. Flexible Intuitions of Euclidean Geometry in an Amazonian Indigene Group. Proceedings of the National Academy of Sciences. 2011; 108(24):9782–9787.

Izard V, Spelke ES. Development of Sensitivity to Geometry in Visual Forms. Human evolution. 2009; 23(3):213– 248.

Jacob G, Pramod RT, Katti H, Arun SP. Qualitative Similarities and Differences in Visual Object Representations Between Brains and Deep Networks. Nature Communications. 2021; 12(1):1872. doi: 10.1038/s41467-021-22078-3.

Jas M, Engemann DA, Bekhti Y, Raimondo F, Gramfort A. Autoreject: Automated Artifact Rejection for MEG and EEG Data. NeuroImage. 2017; 159:417–429. doi: 10.1016/j.neuroimage.2017.06.030.

Jas M, Larson E, Engemann DA, Leppäkangas J, Taulu S, Hämäläinen M, Gramfort A. A Reproducible MEG/EEG Group Study With the MNE Software: Recommendations, Quality Assessments, and Good Practices. Frontiers in Neuroscience. 2018; 12:530. doi: 10.3389/fnins.2018.00530.

Jatoi MA, Kamel N, Malik AS, Faye I. EEG Based Brain Source Localization Comparison of sLORETA and eLORETA. Australasian physical & engineering sciences in medicine. 2014; 37(4):713–721.

Jenkinson M, Bannister P, Brady M, Smith S. Improved Optimization for the Robust and Accurate Linear Registration and Motion Correction of Brain Images. NeuroImage. 2002; 17(2):825–841. doi: 10.1006/nimg.2002.1132.

Kanjlia S, Lane C, Feigenson L, Bedny M. Absence of Visual Experience Modifies the Neural Basis of Numerical Thinking. Proceedings of the National Academy of Sciences. 2016; p. 201524982. doi: 10.1073/pnas.1524982113.

Kanwisher N, Chun MM, McDermott J, Ledden PJ. Functional Imaging of Human Visual Recognition. Cognitive Brain Research. 1996; 5(1-2):55–67.

Kayaert G, Biederman I, Op de Beeck HP, Vogels R. Tuning for Shape Dimensions in Macaque Inferior Temporal Cortex. European Journal of Neuroscience. 2005; 22(1):212–224.

Kayaert G, Biederman I, Vogels R. Shape Tuning in Macaque Inferior Temporal Cortex. Journal of Neuroscience. 2003; 23(7):3016–3027.

Kayaert G, Biederman I, Vogels R. Representation of Regular and Irregular Shapes in Macaque Inferotemporal Cortex. Cerebral Cortex. 2005; 15(9):1308–1321.

Kayaert G, Wagemans J. Infants and Toddlers Show Enlarged Visual Sensitivity to Nonaccidental Compared with Metric Shape Changes. i-Perception. 2010; 1(3):149–158. doi: 10.1068/i0397.

Khosla M, Murty NAR, Kanwisher N. Data-Driven Component Modeling Reveals the Functional Organization of High-Level Visual Cortex. Journal of Vision. 2022; 22(14):4184–4184.

Klein A, Ghosh SS, Bao FS, Giard J, Häme Y, Stavsky E, Lee N, Rossa B, Reuter M, Neto EC, Keshavan A. Mindboggling Morphometry of Human Brains. PLOS Computational Biology. 2017; 13(2):e1005350. doi: 10.1371/jour-nal.pcbi.1005350.

Kourtzi Z, Erb M, Grodd W, Bülthoff HH. Representation of the Perceived 3-D Object Shape in the Human Lateral Occipital Complex. Cerebral cortex. 2003; 13(9):911–920.

Kourtzi Z, Kanwisher N. Cortical Regions Involved in Perceiving Object Shape. Journal of Neuroscience. 2000; 20(9):3310–3318.

Kriegeskorte N, Mur M, Bandettini PA. Representational Similarity Analysis-Connecting the Branches of Systems Neuroscience. Frontiers in systems neuroscience. 2008; 2:4.

Kriegeskorte N, Mur M, Ruff DA, Kiani R, Bodurka J, Esteky H, Tanaka K, Bandettini PA. Matching Categorical Object Representations in Inferior Temporal Cortex of Man and Monkey. Neuron. 2008; 60(6):1126–1141. doi: 10.1016/j.neuron.2008.10.043.

Kubilius J, Schrimpf M, Kar K, Rajalingham R, Hong H, Majaj N, Issa E, Bashivan P, Prescott-Roy J, Schmidt K, Nayebi A, Bear D, Yamins DL, DiCarlo JJ. Brain-Like Object Recognition with High-Performing Shallow Recurrent ANNs. In: Advances in Neural Information Processing Systems, vol. 32 Curran Associates, Inc.; 2019. https://proceedings.neurips.cc/paper/2019/hash/7813d1590d28a7dd372ad54b5d29d033-Abstract.html.

Kubilius J, Schrimpf M, Nayebi A, Bear D, Yamins DLK, DiCarlo JJ, CORnet: Modeling the Neural Mechanisms of Core Object Recognition. Neuroscience; 2018. http://biorxiv.org/lookup/doi/10.1101/408385, doi: 10.1101/408385.

Lake BM, Salakhutdinov R, Tenenbaum JB. Human-level concept learning through probabilistic program induction. Science (New York, NY). 2015; 350(6266):1332–1338. doi: 10.1126/science.aab3050.

Lake BM, Ullman TD, Tenenbaum JB, Gershman SJ. Building Machines That Learn and Think Like People. The Behavioral and Brain Sciences. 2016; p. 1–101. doi: 10.1017/S0140525X16001837.

Lanczos C. Evaluation of Noisy Data. Journal of the Society for Industrial and Applied Mathematics Series B Numerical Analysis. 1964; 1(1):76–85. doi: 10.1137/0701007.

Leeuwenberg ELJ. A Perceptual Coding Language for Visual and Auditory Patterns. The American Journal of Psychology. 1971; 84(3):307. doi: 10.2307/1420464.

Lescroart MD, Biederman I. Cortical Representation of Medial Axis Structure. Cerebral cortex. 2013; 23(3):629–637.

Leyton M. A Generative Theory of Shape, vol. 2145. Springer; 2003.

Liu Z, Mao H, Wu CY, Feichtenhofer C, Darrell T, Xie S. A Convnet for the 2020s. In: Proceedings of the IEEE/CVF conference on computer vision and pattern recognition; 2022. p. 11976–11986.

Long B, Fan JE, Huey H, Chai Z, Frank MC. Parallel Developmental Changes in Children’s Production and Recognition of Line Drawings of Visual Concepts. Nature Communications. 2024; 15(1):1191. doi: 10.1038/s41467-023-44529-9.

Lowet AS, Firestone C, Scholl BJ. Seeing Structure: Shape Skeletons Modulate Perceived Similarity. Attention, Perception, & Psychophysics. 2018; 80:1278–1289.

Lundqvist D, Flykt A, Öhman A. Karolinska Directed Emotional Faces. Cognition and Emotion. 1998;.

Malach R, Reppas J, Benson R, Kwong K, Jiang H, Kennedy W, Ledden P, Brady T, Rosen B, Tootell R. Object-Related Activity Revealed by Functional Magnetic Resonance Imaging in Human Occipital Cortex. Proceedings of the National Academy of Sciences. 1995; 92(18):8135–8139.

Margalit E, Jamison KW, Weiner KS, Vizioli L, Zhang RY, Kay KN, Grill-Spector K. Ultra-High-Resolution fMRI of Human Ventral Temporal Cortex Reveals Differential Representation of Categories and Domains. The Journal of Neuroscience. 2020; 40(15):3008–3024. doi: 10.1523/JNEUROSCI.2106-19.2020.

Mayilvahanan P, Zimmermann RS, Wiedemer T, Rusak E, Juhos A, Bethge M, Brendel W, In Search of Forgotten Domain Generalization. arXiv; 2025. http://arxiv.org/abs/2410.08258, doi: 10.48550/arXiv.2410.08258.

Meyer EE, Martynek M, Kastner S, Livingstone MS, Arcaro MJ. Expansion of a Conserved Architecture Drives the Evolution of the Primate Visual Cortex. Proceedings of the National Academy of Sciences. 2025; 122(3):e2421585122.

Morfoisse T, Izard V. A Unifying Parametric Language and a Toolbox for Future Investigations of the Role of the Medial Axis Parameters in Human Vision. Journal of Vision. 2021; 21(9):2621–2621.

Nishimura M, Scherf KS, Zachariou V, Tarr MJ, Behrmann M. Size Precedes View: Developmental Emergence of Invariant Object Representations in Lateral Occipital Complex. Journal of cognitive neuroscience. 2015; 27(3):474–491.

Niso G, Gorgolewski KJ, Bock E, Brooks TL, Flandin G, Gramfort A, Henson RN, Jas M, Litvak V, T Moreau J, Oostenveld R, Schoffelen JM, Tadel F, Wexler J, Baillet S. MEG-BIDS, the Brain Imaging Data Structure Extended to Magnetoencephalography. Scientific Data. 2018; 5(1):180110. doi: 10.1038/sdata.2018.110.

Oquab M, Darcet T, Moutakanni T, Vo H, Szafraniec M, Khalidov V, Fernandez P, Haziza D, Massa F, El-Nouby A, others. Dinov2: Learning Robust Visual Features Without Supervision. arXiv preprint arXiv:230407193. 2023;.

Papale P, Betta M, Handjaras G, Malfatti G, Cecchetti L, Rampinini A, Pietrini P, Ricciardi E, Turella L, Leo A. Common Spatiotemporal Processing of Visual Features Shapes Object Representation. Scientific Reports. 2019; 9(1):7601.

Papale P, Leo A, Handjaras G, Cecchetti L, Pietrini P, Ricciardi E. Shape Coding in Occipito-Temporal Cortex Relies on Object Silhouette, Curvature, and Medial Axis. Journal of Neurophysiology. 2020; 124(6):1560–1570.

Pascual-Marqui RD, Faber P, Kinoshita T, Kochi K, Milz P, Nishida K, Yoshimura M. Comparing EEG/MEG Neuroimaging Methods Based on Localization Error, False Positive Activity, and False Positive Connectivity. BioRxiv. 2018; p. 269753.

Pernet CR, Appelhoff S, Gorgolewski KJ, Flandin G, Phillips C, Delorme A, Oostenveld R. EEG-BIDS, an Extension to the Brain Imaging Data Structure for Electroencephalography. Scientific Data. 2019; 6(1):103. doi: 10.1038/s41597-019-0104-8.

Pinel P, Dehaene S, Rivière D, LeBihan D. Modulation of Parietal Activation by Semantic Distance in a Number Comparison Task. Neuroimage. 2001; 14(5):1013–1026.

Pinheiro-Chagas P, Daitch A, Parvizi J, Dehaene S. Brain Mechanisms of Arithmetic: A Crucial Role for Ventral Temporal Cortex. Journal of cognitive neuroscience. 2018; 30(12):1757–1772.

Power JD, Mitra A, Laumann TO, Snyder AZ, Schlaggar BL, Petersen SE. Methods to Detect, Characterize, and Remove Motion Artifact in Resting State fMRI. NeuroImage. 2014; 84(Supplement C):320–341. doi: 10.1016/j.neuroimage.2013.08.048.

Quilty-Dunn J, Porot N, Mandelbaum E. The Best Game in Town: The Re-Emergence of the Language of Thought Hypothesis Across the Cognitive Sciences. Behavioral and Brain Sciences. 2022; p. 1–55. doi: 10.1017/S0140525X22002849.

Rauschecker AM, Bowen RF, Parvizi J, Wandell BA. Position Sensitivity in the Visual Word Form Area. Proceedings of the National Academy of Sciences. 2012; 109(24). https://pnas.org/doi/full/10.1073/pnas.1121304109, doi: 10.1073/pnas.1121304109.

Sablé-Meyer M, Ellis K, Tenenbaum J, Dehaene S. A Language of Thought for the Mental Representation of Geometric Shapes. Cognitive Psychology. 2022; 139:101527. doi: 10.1016/j.cogpsych.2022.101527.

Sablé-Meyer M, Fagot J, Caparos S, Kerkoerle Tv, Amalric M, Dehaene S. Sensitivity to Geometric Shape Regularity in Humans and Baboons: A Putative Signature of Human Singularity. Proceedings of the National Academy of Sciences. 2021; 118(16). https://www.pnas.org/content/118/16/e2023123118, doi: 10.1073/pnas.2023123118.

Saito A, Hayashi M, Takeshita H, Matsuzawa T. The Origin of Representational Drawing: A Comparison of Human Children and Chimpanzees. Child Development. 2014; p. n/a–n/a. doi: 10.1111/cdev.12319.

Sassenhagen J, Draschkow D. Cluster-Based Permutation Tests of MEG/EEG Data Do Not Establish Significance of Effect Latency or Location. Psychophysiology. 2019; 56(6):e13335.

Satterthwaite TD, Elliott MA, Gerraty RT, Ruparel K, Loughead J, Calkins ME, Eickhoff SB, Hakonarson H, Gur RC, Gur RE, Wolf DH. An Improved Framework for Confound Regression and Filtering for Control of Motion Artifact in the Preprocessing of Resting-State Functional Connectivity Data. NeuroImage. 2013; 64(1):240–256. doi: 10.1016/j.neuroimage.2012.08.052.

Schrimpf M, Kubilius J, Hong H, Majaj NJ, Rajalingham R, Issa EB, Kar K, Bashivan P, Prescott-Roy J, Geiger F, Schmidt K, Yamins DLK, DiCarlo JJ, Brain-Score: Which Artificial Neural Network for Object Recognition Is Most Brain-Like? Neuroscience; 2018. http://biorxiv.org/lookup/doi/10.1101/407007, doi: 10.1101/407007.

Schuhmann C, Beaumont R, Vencu R, Gordon C, Wightman R, Cherti M, Coombes T, Katta A, Mullis C, Wortsman M, others. Laion-5b: An Open Large-Scale Dataset for Training Next Generation Image-Text Models. Advances in neural information processing systems. 2022; 35:25278–25294.

Sueur C. From Stones to Sketches: Investigating Tracing Behaviours in Japanese Macaques. Primates. 2025; p. 1–6.

Tanaka M. Recognition of Pictorial Representations by Chimpanzees (Pan Troglodytes). Animal cognition. 2007; 10:169–179.

Taulu S, Kajola M. Presentation of Electromagnetic Multichannel Data: The Signal Space Separation Method. Journal of Applied Physics. 2005; 97(12):124905.

Thompson JAF, Sheahan H, Summerfield C. Learning to Count Visual Objects by Combining “What” and “Where” in Recurrent Memory. In: Lourentzou I, Wu J, Kashyap S, Karargyris A, Celi LA, Kawas B, Talathi S, editors. Proceedings of The 1st Gaze Meets ML workshop, vol. 210 of Proceedings of Machine Learning Research PMLR; 2023. p. 199–218. https://proceedings.mlr.press/v210/thompson23a.html.

Tsao DY, Livingstone MS. Mechanisms of Face Perception. Annual Review of Neuroscience. 2008; 31(1):411–437. doi: 10.1146/annurev.neuro.30.051606.094238.

Tsao DY, Moeller S, Freiwald WA. Comparing face patch systems in macaques and humans. Proceedings of the National Academy of Sciences of the United States of America. 2008; 105(49):19514–19519. doi: 10.1073/pnas.0809662105.

Tustison NJ, Avants BB, Cook PA, Zheng Y, Egan A, Yushkevich PA, Gee JC. N4ITK: Improved N3 Bias Correction. IEEE Transactions on Medical Imaging. 2010; 29(6):1310–1320. doi: 10.1109/TMI.2010.2046908.

Tversky B. Visualizing Thought. Topics in Cognitive Science. 2011; 3(3):499–535. doi: 10.1111/j.1756-8765.2010.01113.x.

Uusitalo MA, Ilmoniemi RJ. Signal-Space Projection Method for Separating MEG or EEG into Components. Medical and biological engineering and computing. 1997; 35(2):135–140.

Vinckier F, Dehaene S, Jobert A, Dubus JP, Sigman M, Cohen L. Hierarchical Coding of Letter Strings in the Ventral Stream: Dissecting the Inner Organization of the Visual Word-Form System. Neuron. 2007; 55(1):143–156. doi: 10.1016/j.neuron.2007.05.031.

Van der Waerden BL. Geometry and Algebra in Ancient Civilizations. Springer Science & Business Media; 2012.

Walther A, Nili H, Ejaz N, Alink A, Kriegeskorte N, Diedrichsen J. Reliability of Dissimilarity Measures for Multi-Voxel Pattern Analysis. NeuroImage. 2016; 137:188–200. doi: 10.1016/j.neuroimage.2015.12.012.

Williams MA, Baker CI, Op de Beeck HP, Mok Shim W, Dang S, Triantafyllou C, Kanwisher N. Feedback of Visual Object Information to Foveal Retinotopic Cortex. Nat Neurosci. 2008; 11(12):1439–1445.

Xu Y, Vignali L, Sigismondi F, Crepaldi D, Bottini R, Collignon O. Similar Object Shape Representation Encoded in the Inferolateral Occipitotemporal Cortex of Sighted and Early Blind People. PLOS Biology. 2023; 21(7):e3001930. doi: 10.1371/journal.pbio.3001930.

Xu Y. A Tale of Two Visual Systems: Invariant and Adaptive Visual Information Representations in the Primate Brain. Annual review of vision science. 2018; 4(1):311–336.

Yamins DL, Hong H, Cadieu CF, Solomon EA, Seibert D, DiCarlo JJ. Performance-Optimized Hierarchical Models Predict Neural Responses in Higher Visual Cortex. Proceedings of the National Academy of Sciences. 2014; 111(23):8619–8624.

Zhan M, Goebel R, de Gelder B. Ventral and Dorsal Pathways Relate Differently to Visual Awareness of Body Postures Under Continuous Flash Suppression. eneuro. 2018; 5(1):ENEURO.0285–17.2017. doi: 10.1523/ENEURO.0285-17.2017.

Zhang Y, Brady M, Smith S. Segmentation of Brain MR Images Through a Hidden Markov Random Field Model and the Expectation-Maximization Algorithm. IEEE Transactions on Medical Imaging. 2001; 20(1):45–57. doi: 10.1109/42.906424.

